# Distinct functions of EHMT1 and EHMT2 in cancer chemotherapy and immunotherapy

**DOI:** 10.1101/2023.10.03.560719

**Authors:** Zhihua Kang, Pan Fu, Hui Ma, Tao Li, Kevin Lu, Juan Liu, Vasudeva Ginjala, Peter Romanienko, Zhaohui Feng, Ming Guan, Shridar Ganesan, Bing Xia

## Abstract

EHTM1 (GLP) and EHMT2 (G9a) are closely related protein lysine methyltransferases often thought to function together as a heterodimer to methylate histone H3 and non-histone substrates in diverse cellular processes including transcriptional regulation, genome methylation, and DNA repair. Here we show that EHMT1/2 inhibitors cause ATM-mediated slowdown of replication fork progression, accumulation of single-stranded replication gaps, emergence of cytosolic DNA, and increased expression of STING. EHMT1/2 inhibition strongly potentiates the efficacy of alkylating chemotherapy and anti-PD-1 immunotherapy in mouse models of tripe negative breast cancer. The effects on DNA replication and alkylating agent sensitivity are largely caused by the loss of EHMT1-mediated methylation of LIG1, whereas the elevated STING expression and remarkable response to immunotherapy appear mainly elicited by the loss of EHMT2 activity. Depletion of UHRF1, a protein known to be associated with EHMT1/2 and LIG1, also induces STING expression, and depletion of either EHMT2 or UHRF1 leads to demethylation of specific CpG sites in the *STING1* promoter, suggestive of a distinct EHMT2-UHRF1 axis that regulates DNA methylation and gene transcription. These results highlight distinct functions of the two EHMT paralogs and provide enlightening paradigms and corresponding molecular basis for combination therapies involving alkylating agents and immune checkpoint inhibitors.

## Introduction

EHMT1 (euchromatic histone methyltransferase I, also known as GLP) and its paralog EHMT2 (G9a) belong to the SET domain proteins that methylate lysine residues such as histone H3K9^1–3^. Methylated H3K9 serves as a chromatin mark for transcriptional inactivation to suppress the expression of a plethora of genes involved in diverse cellular functions, including embryonic development, cell proliferation, metabolism, and other biological processes^4–6^. Non-histone proteins including p53, HIF1α, and DNMT3A have also been reported to be methylated by EHMT1 and EHMT2, but the biological significance of these protein methylation events remains largely elusive^7–10^. The two orthologs form in a heterodimeric complex and are often thought to function together.

EHMT1/2 associate with PCNA on the replication fork^11^ and methylate LIG1 (DNA ligase I), the main ligase for Okazaki fragments, on K126, and this modification enables LIG1 to interact with UHRF1 (ubiquitin like with PHD and ring finger domains 1)^12^. UHRF1 is known to be required for DNMT1 recruitment to DNA methylation sites^13^, and K126-methylated LIG1 may recruit DNMT1 through UHRF1 to the Okazaki fragments to methylate newly synthesized lagging strands to maintain genome methylation patterns^12^. Very recently, EHMT2 is reported to catalyze heterochromatin assembly at stressed replication forks to ensure replication fork stability^14^. However, whether the EHMT1/2-LIG1-UHRF1 axis regulates DNA replication or repair per se is unclear. Moreover, we and others have shown that EHMT1 and EHMT2 are recruited to DNA damage sites promote DNA repair^15–17^.

EHMT1/2 inhibition has been reported to inhibit the proliferation of cancer cells and enhances cancer cell killing by various chemotherapeutic agents^18^. Recently, inhibition of EHMT2 has also been reported to alter the immune microenvironment of melanoma^19^, bladder cancer^20^, and ovarian serous carcinoma^21^, which enhances the anti-tumor effect of immune checkpoint inhibitors (ICIs), yet the mechanisms and key mediators in this process are also unclear.

In the present manuscript, we demonstrate the distinct roles of EHMT1 and EHMT2 in DNA replication, DNA repair and DNA methylation. In particular, we show that inhibition of EHMT1-mediated methylation of LIG1 causes a slowdown of replication fork progression and sensitizes cancer cells to alkylating agents, whereas inhibition of EHMT2 induces the expression of STING (stimulator of interferon genes) and promotes tumor response to ICIs.

## Results

### EHMT1 inhibition induces replication fork slowdown

To explore the function of EHMT1/2 in DNA replication, we treated U2OS (osteosarcoma) cells with two EHMT1/2 inhibitors, UNC0638 and UNC0642^22, 23^, and analyzed their impact on a panel of replication factors and several DNA replication-related parameters. Interestingly, levels of chromatin bound LIG1 were substantially reduced in the inhibitors treated cells, although the change of total LIG1 expression was not consistent in different repeats (Fig. 1a, Extended Data Fig. 1a). In contrast, an increase in chromatin bound LIG3 levels were observed, suggestive of a possible compensation for the reduced LIG1. Total UHRF1 levels were moderately lower in the drug-treated cells, and its levels in the chromatin fraction were also slightly reduced. Little to no changes were observed for other replication factors such as MCM4, MCM10, CDC45, and PCNA.

**Figure 1.**
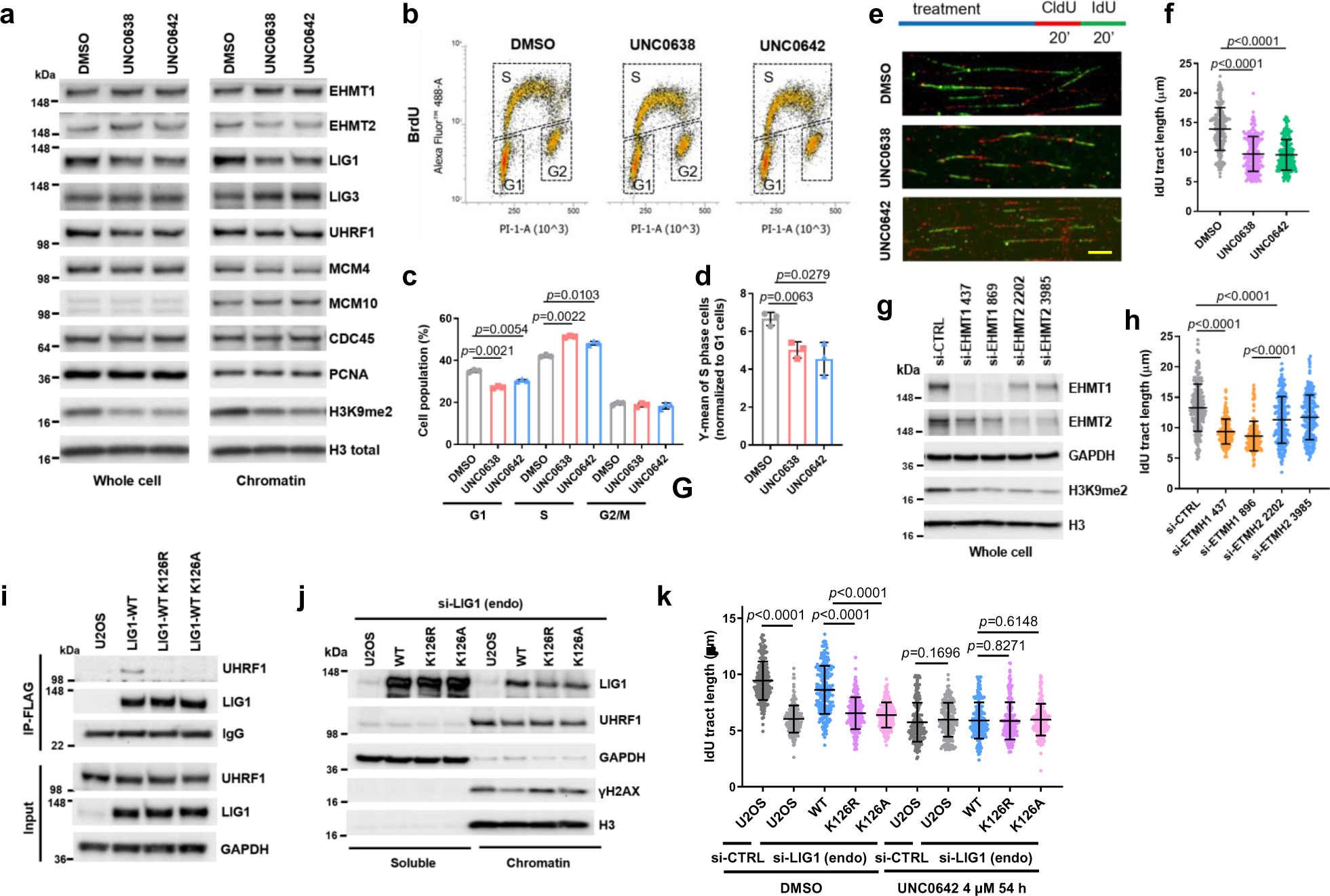
EHMT1/EHMT2 inhibition or EHMT1 depletion induces replication stress. **a** Protein levels of DNA replication factors in U2OS cells treated with UNC0638 and UNC0642 (4 µM for 54 h). **b** BrdU incorporation in U2OS cells treated with UNC0638/0642 (4 µM for 54 h). **c** Quantification of cell cycle distributions of cells treated as in **b**. **d** Quantification of BrdU levels in S phase cells after UNC0638/0642 treatments. The value for each S phase population was normalized against G1 population from each condition. **e** Labeling scheme and the representative images of the DNA fiber assay. Scale bar, 10 µm. **f** Lengths of IdU labeled replication tracts in U2OS cells treated as in **a**. **g** siRNA-mediated depletion of EHMT1 or EHMT2 in U2OS cells. **h** Replication tract lengths in EHMT1- or EHMT2-depleted U2OS cells. **i** Co-IP of endogenous UHRF1 with FLAG-HA-tagged WT, K126R and K126A LIG1 in U2OS cells. **j** Protein levels of LIG1, UHRF1 and γH2AX in soluble and chromatin fractions of blank U2OS cells and cells selectively expressing exogenous (WT, K126R and K126A) LIG1 proteins. **k** Lengths of IdU labeled replication tracts in U2OS cells selectively expressing LIG1-WT, K126R or K126A treated with UNC0642 (4 µM for 54 h). Data (**c**, **d**, dots=3; **f**, **h**, **k**, dots=200) are presented as mean ± standard deviation (SD). Statistical test is preformed using two-tailed paired Student’s *t* test in **c**, **d** and unpaired Student’s *t* test in **f**, **h**, **k**.

Cell cycle analysis using the BrdU incorporation assay revealed that the UNC compounds led to an increase in S phase population and a concomitant decrease in G1 population, whereas the G2 population remained largely unchanged (Fig. 1b and c). Interestingly, the intensity of BrdU incorporation appeared to be decreased in the drug-treated cells (Fig. 1d). These results suggest that the rate of DNA replication may be slower upon EHMT1/2 inhibition. Indeed, DNA fiber analysis showed that replication fork speed in the drug-treated cells were significantly slower (Fig. 1e and f). Similar results were also obtained in the MDA-MB-231 triple negative breast cancer (TNBC) cells and the 4T1 mouse mammary tumor cells after EHMT1/2 inhibition (Extended Data Fig. 1b-e).

Next, we asked whether EHMT1 or EHMT2 inhibition is primarily responsible for the effects observed. We found that siRNA-mediated depletion of EHMT1 had a significantly stronger impact on fork progression than EHMT2 depletion (Fig. 1g and h). Note that the two proteins appeared to be very stable, and the depletions were incomplete. Moreover, the two proteins form a complex, and loss of one destabilizes the other. Therefore, based on these results alone one cannot conclude that EHMT1 is more important for DNA replication than EHMT2. However, our observations are consistent with the previously reported finding that *Ehmt1*^−/−^ mouse embryonic stem cells (ES cells) showed much stronger reduction in LIG1-K126 methylation than *Ehmt2*^−/−^ ES cells^12^. Moreover, we also found much less LIG1 in the chromatin fraction of EHMT1-depleted cells compared to EHMT2-depleted cells (Extended Data Fig. 1f). Together, these findings strongly suggest that replication stress elicited by UNC0638 and UNC0642 is mainly mediated by EHMT1 inhibition.

### EHMT1-mediated methylation of LIG1 is required for normal replication fork progression

Given the decrease in LIG1 and (to a lesser extent) UHRF1 abundance in chromatin compartment of cells treated with the EHMT1/2 inhibitors, we tested the impact of LIG and UHRF1 downregulation on replication fork progression. siRNA knockdown of LIG1 led to a substantial reduction in fork speed (Extended Data Fig. 2a and b), while the effect of UHRF1 knockdown was even stronger (Extended Data Fig. 2c and d). Moreover, depletion of either protein led to a clear increase in Ser139-phosphorylated H2AX (γH2AX), a marker of DNA double strand breaks (DSBs), suggestive of replication fork collapse and/or a failure to repair DSBs elsewhere.

As K126 methylation of LIG1 is required for its interaction with UHRF1^12^, we hypothesized that the reduced replication fork progression may result from a loss of LIG-UHRF1 complex formation. To test this hypothesis, we generated stable U2OS cell lines expressing wild-type (WT) or K126R or K126A mutant forms of LIG1. Treatment of cells expressing WT LIG1 with either UNC0638 or UNC0642 led to a loss of UHRF1 association with LIG1 (Extended Data Fig. 2e), and both K126R and K126A mutant failed to bind UHRF1 (Fig. 1i), confirming the published result^12^. In these cells, the expression levels of both mutant LIG1 proteins were similar to that of the WT protein but their abundance in the chromatin fraction was lower (Fig. 1j). A similar observation was also made in 293T cells transiently expressing the WT and mutant LIG proteins (Extended Data Fig. 2f). DNA fiber assay revealed that while exogenous WT LIG1 was able to rescue the reduced fork progression caused by a depletion of the endogenous protein, both K126R and K126A mutants failed to do so (Fig. 1k). Moreover, UNC0642 treatment eliminated the difference between cells expressing WT and mutant LIG1, as all cells showed an equally reduced fork speed (Fig. 1k).

Collectively, the above results indicate that EHMT1-mediated methylation of LIG1 on K126 promotes its chromatin association and replication fork progression, and they further suggest that LIG-UHRF1 complex formation is required for the normal progression of replication forks. Still, it is possible that LIG1 methylation itself also plays a role in fork progression independent of its effect on UHRF1 binding.

### Fork slowdown upon LIG1 depletion or EHMT1 inhibition is mediated by ATM

Given that LIG1-depleted cells showed a significant increase in γH2AX (Extended Data Fig. 2a), we asked whether ATM or ATR, the two apical DNA damage signaling kinases that phosphorylate H2AX, is responsible for triggering the fork slowing in the cells. Indeed, LIG1 depletion in U2OS cells led to ATM activation, as evidenced by increased phosphorylation of its substrates KAP1 S824 and CHK2 T68 (Fig. 2a), and inhibition of ATM with a small molecule inhibitor KU55933 fully rescued the fork slowdown caused by LIG1 depletion (Fig. 2b). When the cells were treated with ionizing radiation (IR) to induce further DNA damage, replication forks slowed further, and ATM inhibition still led to major rescue of fork progression, especially in LIG-depleted cells (Fig. 2b). In contrast, ATR inhibition with VE821 led to a further slowdown, rather than a rescue, of fork progression (Fig. 2c).

**Figure 2.**
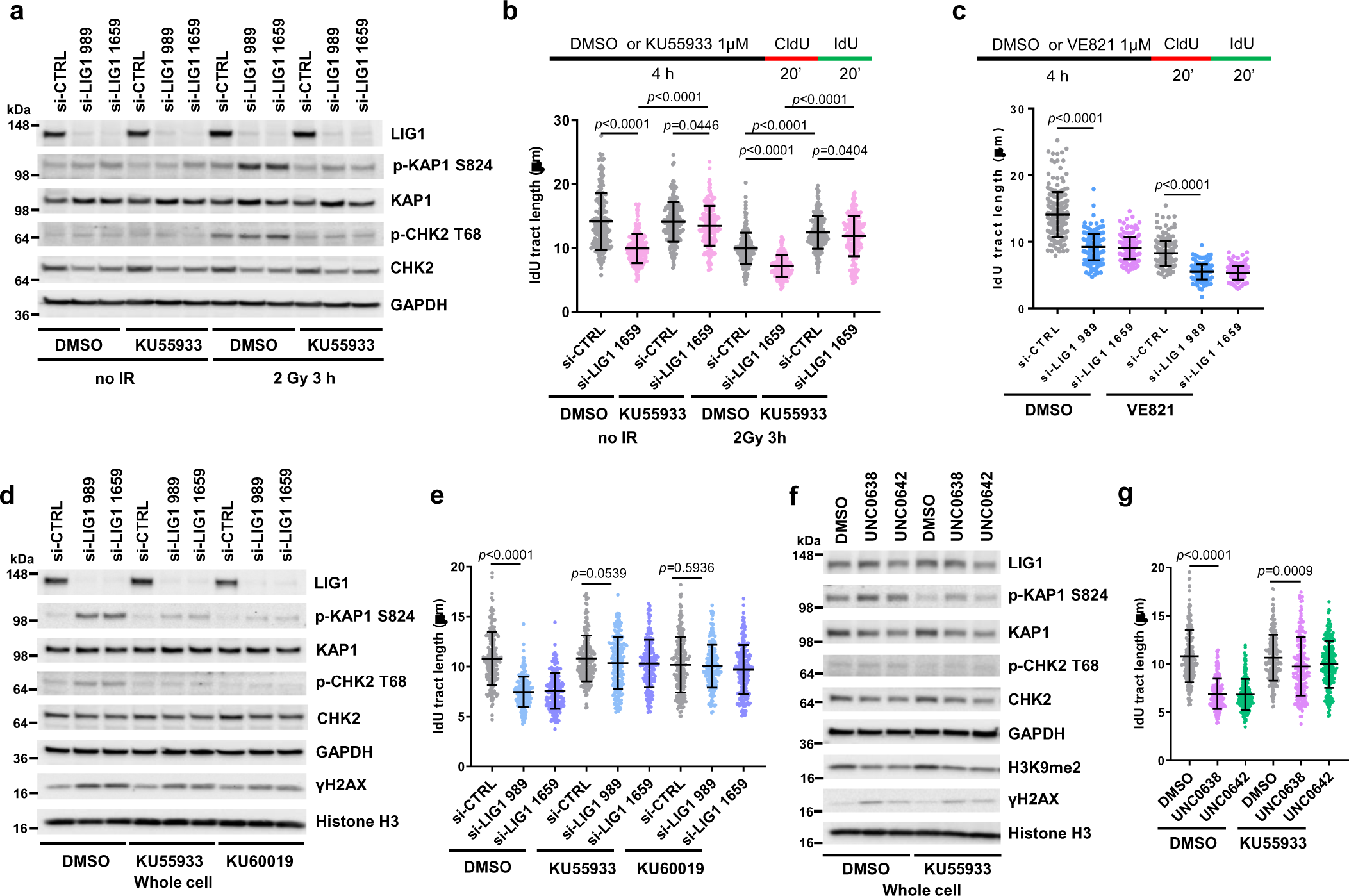
EHMT1 inhibition restrain replication fork progression through ATM activation. **a** Western blots showing levels of KAP1, CHK2 and H2AX phosphorylation in U2OS cells treated as indicated. Duplicate set of cells were treated with siRNAs for 72 h followed by DMSO or KU55933 for 1 h, and then one set of cells were subjected to 2 Gy of IR and the other unirradiated. Cells were collected 3 h post IR. **b** Lengths of IdU labeled replication tracts in LIG1-depleted U2OS cells treated with DMSO or KU55933 without or with IR treatment. **c** Lengths of IdU labeled replication tracts in LIG1-depleted U2OS cells treated with DMSO or ATRi VE821. **d** Western blots showing levels of KAP1, CHK2 and H2AX phosphorylation in MDA-MB-231 cells treated with control or LIG1 siRNAs for 72 h and then with DMSO, KU55933 (1 µM), or KU60019 (0.4 µM) for 4 h. **e** Lengths of IdU labeled replication tracts in MDA-MB-231 cells treated as in **d**. **f** Western blots showing levels of KAP1, CHK2 and H2AX phosphorylation in MDA-MB-231 cells treated with DMSO, UNC0638, or UNC0642 (4 µM for 54 h) and then with DMSO, KU55933 (1 µM) for 4 h. **g** Lengths of IdU labeled replication tracts in MDA-MB-231 cells treated as in **f**. Data (**b**, **c**, **e**, **g** dots=200) are presented as mean ± standard deviations (SD). Statistical test is preformed using two-tailed unpaired Student’s *t* test and the exact *p* value is provided.

We next evaluated the effect of LIG1 depletion in the MDA-MB-231 breast cancer cells. In these cells, knockdown of LIG1 also induced KAP1 and CHK2 phosphorylation, indicative of ATM activation, and the elevated phosphorylation levels were reversed by two different ATM inhibitors, KU55933 and KU60019 (Fig. 2d). The downregulation of replication fork elongation caused by LIG1 knockdown was also rescued by the two inhibitors, to a level similar to that in control siRNA treated cells (Fig. 2e).

Finally, the impact of EHMT1/2 inhibition by UNC0638 and UNC0642 in MDA-MB-231 cells was tested. Similar to LIG1 depletion, the two compounds each induced KAP1 and CHK2 phosphorylation that could be reversed by ATM inhibition (Fig. 2f). DNA fiber analysis also showed that the decreased replication fork progression caused by the compounds was rescued after ATM inhibition (Fig. 2g). In light of the previous report that CHK2 phosphorylates components of the CDC45-MCM-GINS (CMG) complex and inhibits its activity^24^, our results suggest that EHMT1 inhibition induced impairment of LIG1 function results in nicks in DNA during replication and repair ultimately leading to DSB formation and ATM-CHK2 activation, which plays a major role in slowing down the replication fork progression.

### LIG1 loss or EHMT1 inhibition selectively sensitizes cancer cells to alkylating agents

To better understand the role of LIG1 in different DNA repair processes, we measured the sensitivity of LIG1-depleted MDA-MB-231 cells to several DNA damaging chemotherapeutic agents, including bleomycin (BLEO, a radiomimetic), camptothecin (CPT, a topoisomerase I toxin), cisplatin (CDDP, a crosslinker), and cyclophosphamide (CTX, an alkylating agent). While depletion of LIG1 caused little to no effect on cellular sensitivity to BLEO, CPT and CDDP, it significantly sensitized cells to CTX (Fig. 3a). Note that CTX, which is used to treat breast, ovarian, lymphoid and other cancers, is a prodrug that is metabolized and converted into its active form in the liver, therefore a high concentration is commonly used in tissue culture studies, while much lower doses are used *in vivo*.

**Figure 3.**
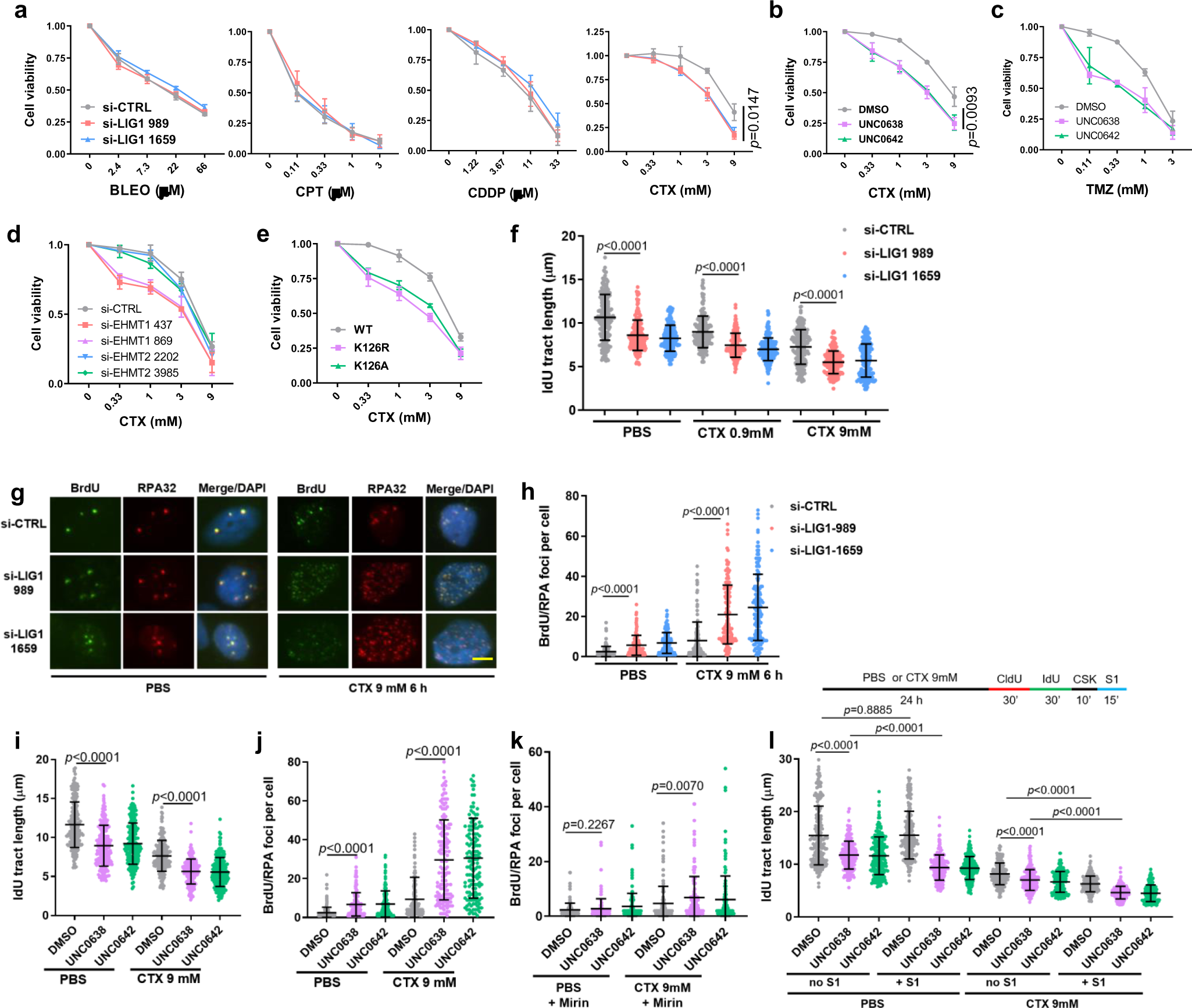
EHMT1 inhibition sensitizes cancer cells to cyclophosphamide through enhancing DNA damage accumulation and replication stress. **a** Survival curves of LIG1-depleted MDA-MB-231 cells treated with different concentrations of bleomycin (BLEO), camptothecin (CPT), cisplatin (CDDP), or cyclophosphamide (CTX) for 72 h. **b-c** Survival curves of MDA-MB-231 cells treated with different concentrations of CTX (b) or temozolomide (TMZ, c) in combination with a fixed concentration (2 µM) of UNC0638 or UNC0642 for 72 h. **d** Survival curves of EHMT1 or EHMT2 depleted MDA-MB-231 cells treated with different concentrations CTX of for 72 h. **e** Survival curves of U2OS cells selectively expressing exogenous LIG1-WT, K126R or K126A following exposure to different concentrations of CTX for 72 h. **f** Lengths of IdU labeled replication tracts in MDA-MB-231 cells treated with LIG1 siRNAs for 48 h and then CTX for 24 h **g** IF images showing BrdU and RPA foci in control and LIG1-depleted MDA-MB-231 cells treated with CTX. Scale bar, 5 µm. **h** Quantification of colocalized BrdU/RPA foci in MDA-MB-231 cells treated as in e. **i** Lengths of IdU labeled replication tracts in MDA-MB-231 cells treated with DMSO or UNC0638/0642 (4 µM) for 30 h and then CTX for 24 h. **j** Numbers of colocalized BrdU/RPA foci in MDA-MB-231 cells treated with DMSO or UNC0638/0642 (4 µM) for 48 h and then CTX for 6 h. **k** Numbers of colocalized BrdU/RPA foci in MDA-MB-231 cells treated as in j and then incubated with mirin. **l** Lengths of IdU labeled replication tracts in MDA-MB-231 cells treated as in i and then incubated with S1 endonuclease. Data (**f**, **i**, **l**, dots=200; **h**, **j**, **k**, dots=150) are presented as mean ± standard deviation (SD). Statistical test is performed using two-tailed unpaired Student’s *t* test.

Considering that UNC0638 and UNC0642 cause reduced LIG1 chromatin association and loss of LIG1-UHRF1 complex formation, we tested the efficacy of their combination with CTX. As the two drugs cause cell death by themselves at higher concentrations (Extended Data Fig. 3a), we used a relatively low concentration (2 µM) of these two inhibitors in the combination. Indeed, both agents significantly enhanced the killing of MDA-MB-231 cells by CTX (Fig. 3b). We also tested combinations of the UNC compounds with temozolomide (TMZ), another alkylating chemotherapeutic agent commonly used to treat brain cancers such as glioblastoma (GBM), and again found a clear enhancement in cell killing (Fig. 3c). Enhancement of cell killing of CTX by the UNC compounds were also seen in U2OS and 4T1 cells (Extended Data Fig. 3b, c).

We then compared the combination of either EHMT1 or EHMT2 depletion and CTX in cancer cell killing and found that depletion of EHMT1, and not EHMT2, sensitized MDA-MB-231 cells to CTX (Fig. 3d), confirming that the effect of UNC0638 and UNC0642 was likely mediated by EHMT1 rather than EHMT2 inhibition. To address whether the effect of the UNC compounds is due to loss of LIG1 methylation, we depleted the endogenous LIG1 in the afore-mentioned U2OS cells stably expressing WT, K126A and K126R LIG1 and then treated the cells with CTX, and we found that cells selectively expressing either mutant protein were hypersensitive to the agent (Fig. 3e). These results clearly demonstrate that the compounds sensitize cancer cells to alkylating agents by inhibiting EHMT1-mediated LIG1 methylation at K126.

We next explored how loss of LIG1 sensitizes cancer cells to CTX. Depletion of LIG1 caused DNA damage and ATM activation, while combination of LIG1 depletion and CTX treatment led to further elevated DNA damage and ATM activation (Extended Data Fig. 3d). Moreover, the combination also caused further slowdown of replication fork progression than either LIG1 depletion or CTX treatment alone (Fig. 3f). We subsequently labeled DNA in MDA-MB-231 cells with BrdU for an extended period of time and then used immunofluorescence (IF) to assess the levels of single-stranded DNA (ssDNA) in the cells. Under non-denaturing conditions, appearance of BrdU foci would reflect long stretches of ssDNA, and co-localization of BrdU with RPA32, a commonly used marker of ssDNA, would further confirm the indication. We found that depletion of LIG1 or CTX treatment each caused a moderate increase in ssDNA, whereas treatment of LIG1-depleted cells with CTX led to a much more dramatic increase in a large fraction of cells (Fig. 3g and h). Such excessive ssDNA may cause RPA exhaustion that severely impairs essential DNA metabolism thereby contributing to cell killing.

Similar to LIG1 depletion, combination of either UNC0638 or UNC0642 with CTX led to further elevated DNA damage and ATM activation (Extended Data Fig. 3e), as well as further decreased replication fork progression (Fig. 3i), accompanied by further increased ssDNA accumulation as revealed by co-localized BrdU and RPA32 foci (Fig. 3j). We then asked whether a nuclease is responsible for the ssDNA generation. A prime candidate is MRE11, which plays key roles in DNA repair and replication fork stability control^25^. Indeed, an inhibitor of MRE11 exonuclease activity, mirin, largely reversed the dramatic increase in ssDNA in MDA-MB-231 cells treated with the drugs, especially the combination (compare Fig. 3k with 3j).

As regions of ssDNA can emerge either from various DNA damage repair processes or from gaps in newly replicated DNA, we employed a modified DNA fiber assay^26^ to assess the levels of replication gaps in MDA-MB-231 cells treated with the different compounds or combinations. In this assay, an S1 nuclease treatment step is added so that newly synthesized DNA with single stranded gaps would be cleaved resulting in replication tracts appearing shorter. Interestingly, while control (DMSO) treated cells showed no detectable replication gaps, UNC0638 or UNC0642 treatment alone already led to a significant amount of such gaps (Fig. 3l). CTX alone also caused replication gap formation, and combinations of either UNC0638 or UNC0642 with CTX led to even more gap formation.

Collectively, the above results demonstrate that LIG1 is critical for the repair of alkylated DNA and suggest that alkylating agents may be a targeted therapy for LIG1 low or deficient tumors. Our data also demonstrates that EHMT1 inhibition sensitizes cancer cells to alkylating agents presumably by compromising LIG1 function leading to excessive ssDNA formation, severe replication stress, and DNA breakage (Extended Data Fig. 4).

### EHMT1 inhibition enhances the efficacy of alkylating chemotherapy

To evaluate the potential therapeutic efficacy of EHMT1 inhibition and CTX combination for cancer treatment, we carried out preclinical studies using two different mouse models of TNBC. In the first model, we injected MDA-MB-231 cells into the largely immunodeficient BALB/c nude mice and treated the resulting tumors with UNC0642 and CTX, either alone or in combination, for 7 consecutive days (Fig. 4a). At the dose used (5 mg/kg), UNC0642 showed no effect on the xenograft tumors, while tumor growth in mice treated with CTX (20 mg/kg, a relatively low dose) continued during the first few days of the treatment and then nearly stopped for about 10 days, before gradually resuming; in contrast, tumors treated with UNC0642+CTX combination stopped growing immediately upon treatment and remained almost completely static over the next 3 weeks until the study reached its endpoint (Fig. 4b and c). Immunohistochemistry (IHC) was conducted to assess tumor proliferation (Ki67), cell death (cleaved caspase3) and DNA damage (γH2AX). As shown in Fig. 4d, tumors from mice treated with UNC0642+CTX displayed reduced proliferation, increased DNA damage, and increased apoptosis, consistent with their reduced growth rates.

**Figure 4.**
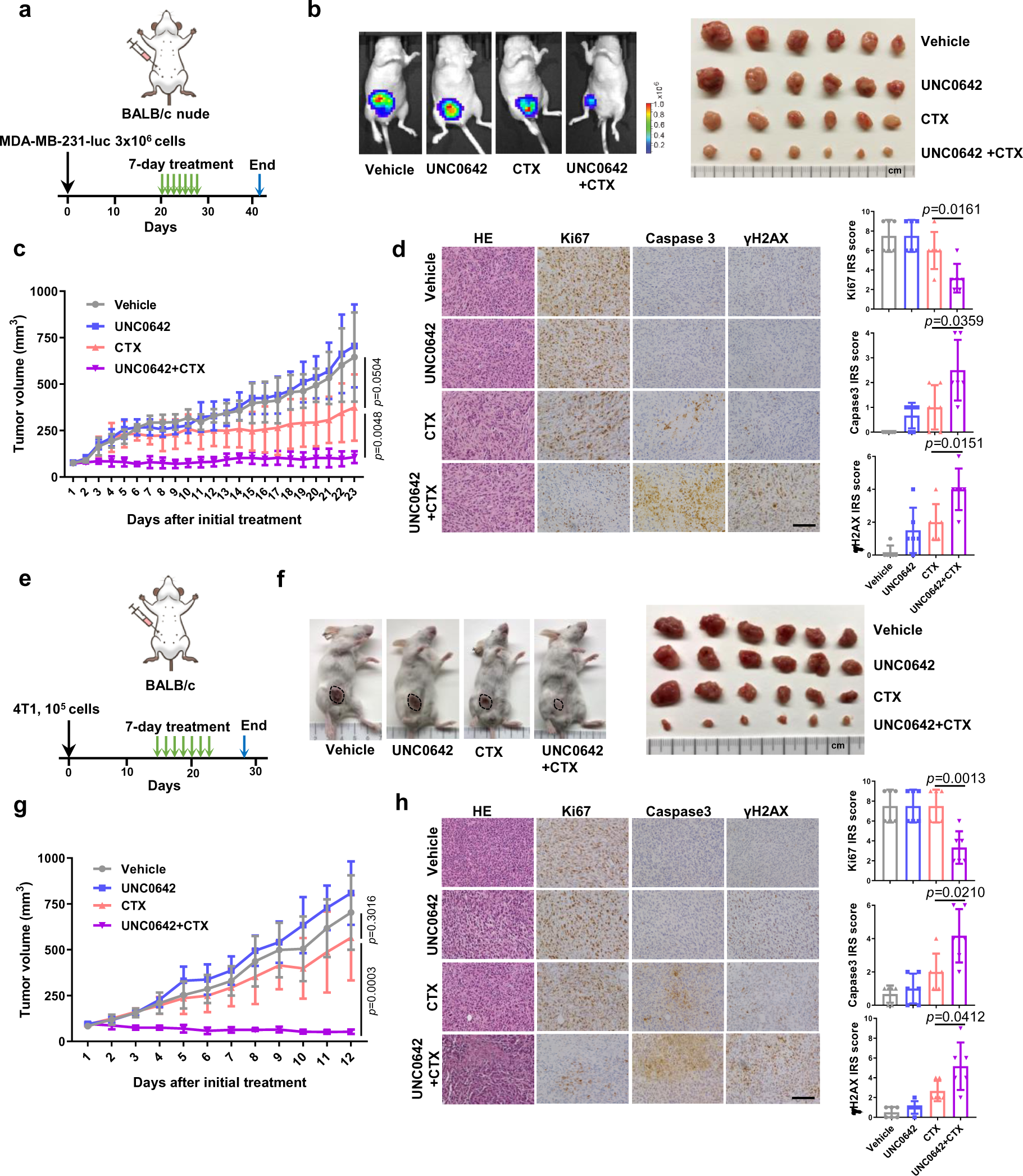
EHMT1 inhibition enhances antitumor response of cyclophosphamide in two triple negative breast cancer mouse models. **a** Schematic diagram of the MDA-MB-231-luc xenograft model and experimental timeline. **b** Representative bioluminescent images of MDA-MB-231-luc tumor bearing mice (left) and an image of all the mammary tumors harvested (right) at study endpoint (day 42). **c** Growth curves of MDA-MB-231-luc tumors in BALB/c nude mice from the first day of treatment (day 20) to study endpoint (day 42). **d** Representative H&E and IHC images of MDA-MB-231-luc tumors (left) and quantification of the IHC results using the immunoreactive score (IRS, right). Scale bar, 50 µm. **e** Schematic diagram of 4T1 allograft models and experimental timeline. **f** Representative images of 4T1 tumor bearing mice (left) and an image of all the mammary tumors harvested at study endpoint (day 25). **g** Growth curves of 4T1 tumors from the first day of treatment (day 14) to study endpoint (day 25). **h** Representative IHC images of 4T1 tumors (left) and quantification of the IHC results using the immunoreactive score (right). Scale bar, 50 µm. Data (**c**, **g**, n=6; **d**, **h**, dots=6) are presented as mean ± SD. Statistical test is preformed using two-tailed unpaired Student’s *t* test and the exact *p* value is provided.

In the second model, we injected the 4T1 mouse mammary tumor cells into normal, immunoproficient BALB/c mice to generate syngeneic allograft tumors and then treated the mice in the same way as in the above xenograft model (Fig. 4e). Again, UNC0642 at the dose used was completely unable to inhibit tumor growth, and CTX alone also failed to produce any significant effect; remarkably, however, combination of the two agents led to tumor shrinkage (Fig. 4f and g). IHC analyses of the tumors again showed reduced proliferation, increased DNA damage, and increased apoptosis (Fig. 4h). Taken together, these data demonstrate that EHMT1 inhibition can substantially enhance the antitumor efficacy of CTX *in vivo*. In fact, in our preliminary studies, tumor disappeared in a mouse upon the combination treatment. Moreover, mice treated with the combination showed little to no weight loss observed following 7 consecutive days of treatment.

### EHMT2 inhibition induces STING expression and activates the cGAS-STING pathway

While examining the histology of the above 4T1 tumors, we noticed increased lymphocytes in the UNC0642-treated group compared with the vehicle-treated group. Indeed, IHC analysis confirmed much more prominent CD4+ and CD8+ T lymphocytes infiltration in the UNC0642 group (Fig. 5a). Considering that UNC0642 treatment led to DNA damage in cells, we tested whether the cGAS-STING pathway^27^, which is activated by cytosolic DNA and known to induce immune response and lymphocytes infiltration, was activated in UNC0642-treated tumors. Interestingly, STING expression level was increased in these tumors (Fig. 5b). Downstream in the pathway, programmed death-ligand 1 (PD-L1) expression was also increased (Fig. 5c), which likely promoted cancer cell immune evasion, and this may explain, at least in part, why even high levels of CD4+ and CD8+ T cells infiltration did not suppress tumor growth (in fact, UNC0642-treated tumors grew slightly faster).

**Figure 5.**
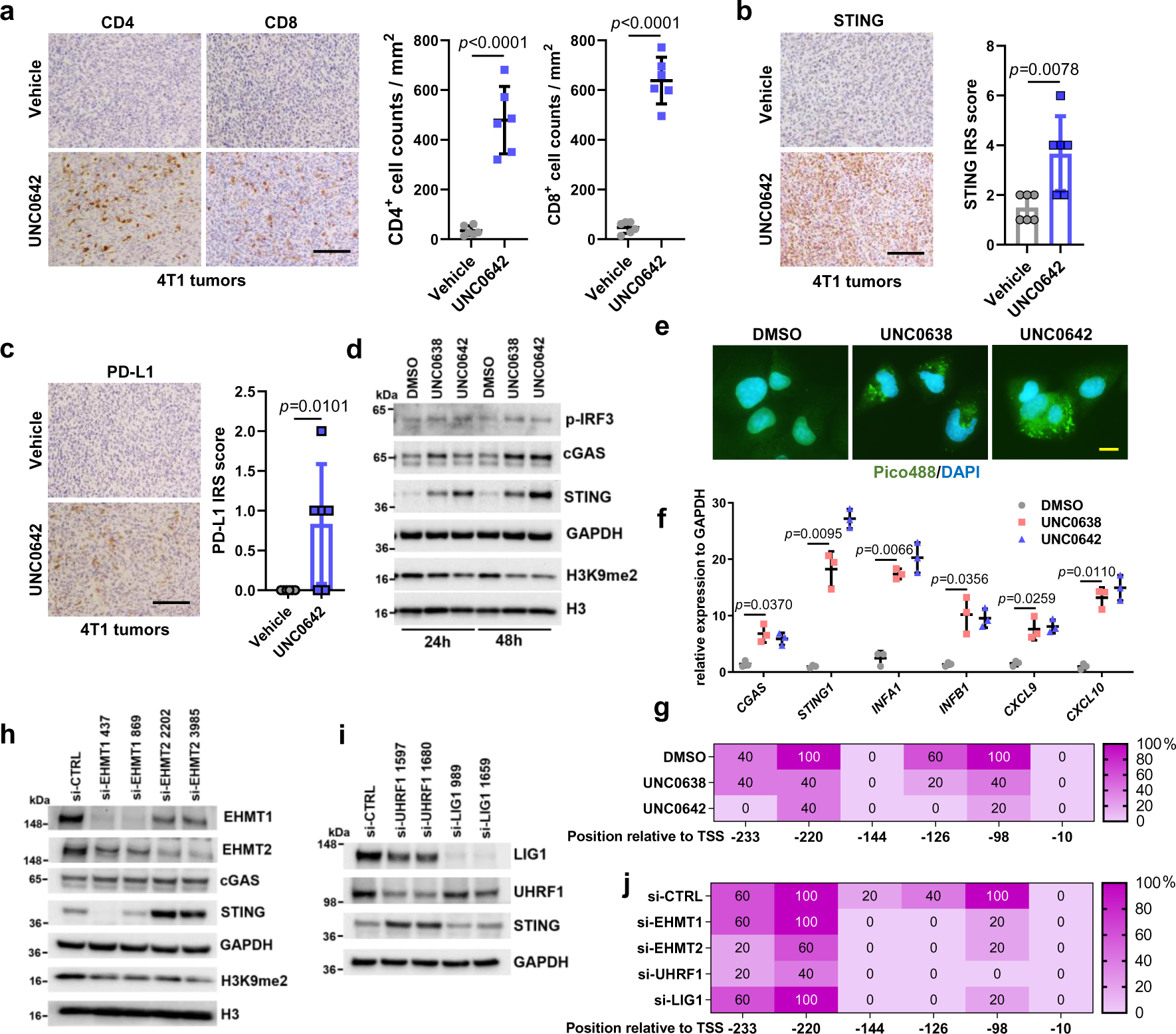
EHMT1/EHMT2 inhibition activates STING signaling and induces lymphocytes infiltration in 4T1 syngeneic allograft tumors. **a** Representative CD4+ and CD8+ staining IHC images of 4T1 tumors (left) and quantification of the infiltrating lymphocytes (right). Scale bar, 50 µm. **b** Representative STING IHC images of 4T1 tumors (left) and quantification using the immunoreactive score (right). Scale bar, 50 µm. **c** Representative PD-L1 IHC images of 4T1 tumors (left) and quantification of the IHC results using the immunoreactive score (right). Scale bar, 50 µm. **d** Protein levels of cGAS-STING pathway factors in U2OS cells treated with DMSO, UNC0638 or UNC0642 (4 µM for 24 and 48 h). **e** Detection of cytosolic DNA using the Pico488 fluorescence dye in U2OS cells treated with DMSO, UNC0638 or UNC0642 (4 µM for 48 h). Scale bar, 10 µm. **f** Quantitative real time (qRT)-PCR analysis (n=3) mRNA expression levels of cGAS-STING pathway genes in U2OS cells treated as in **e**. **g** Percentage of methylation for each CpG site in the *STING1* promoter as determined by sequencing of bisulfite-converted DNA from U2OS cells treated as in **e**. **h** Western blots showing levels of indicated proteins in U2OS cells treated with EHMT1 or EHMT2 siRNAs for 72 h. **i** Western blots showing levels of indicated proteins in U2OS cells treated with LIG1 or UHRF1 siRNAs for 72 h. **j** Percentage of methylation for each CpG site in the *STING1* promoter from U2OS cells treated with EHMT1, EHMT2, UHRF1 and LIG1 siRNAs. Data (**a**, **b**, **c** dots=6; **f**, dots=3) are presented as mean ± standard deviation (SD). Statistical test is preformed using two-tailed unpaired Student’s *t* test in **a**, **b**, **c** and paired Student’s *t* test in **f** and the exact *p* value is provided.

To investigate the potential mechanisms of increased STING expression induced by UNC0642, we first treated U2OS cells with UNC0642 (and UNC0638) and examined the levels of STING and its activator cGAS. The result showed that not only STING, cGAS amount also increased (Fig. 5d). Accordingly, the downstream effector phospo-IRF3 was also elevated. As cGAS, and therefore STING, is activated by cytosolic DNA, we assessed its levels in cells treated with the UNC compounds using the Pico488 fluorescence dye. Indeed, compared to the control (DMSO treated) cells, UNC06238 and UNC0642 treatment led to strong staining of DNA in the cytosol (Fig. 5e). We then performed real-time PCR to evaluate the mRNA levels of *CGAS*, *STING1*, and several downstream genes in the pathway, including interferon genes *INFA1* and *INFB1* and chemokine genes *CXCL9* and *CXCL10*^28^. Remarkable upregulation was seen for all these genes in cells treated with UNC0642 and UNC0638 (Fig. 5f), confirming the activation of the cGAS-STING pathway in the cells.

*STING1* expression has been reported to be regulated by methylation of CpG sites in its promoter region^29, 30^, especially within 250 nt upstream of the transcriptional start site (TSS) (Extended Data Fig. 5a). Through bisulfite sequencing, we determined that both two inhibitors substantially reduced the methylation levels of three CpG sites, at −98, −126 and −220 nt from the TSS, respectively (Fig. 5g, Extended Data Fig. 5b), suggesting that these sites may play a key role in the regulation of *STING1* transcription. In MDA-MB-231 cells, STING expression was also induced by the UNC compounds, and a similar cGAS-STING pathway activation was observed (Extended Data Fig. 5c and d).

To test whether EHMT1, EHMT2, UHRF1 and LIG1 were involved in the regulation of CpG methylation of *STING1*, we knocked down each of these proteins in U2OS cells. Depletion of either EHMT2 or UHRF1, but neither EHMT1 nor LIG1, led to increased STING expression (Fig. 5h and i). Interestingly, bisulfite sequencing revealed that knockdown of each of these four proteins reduced the methylation level at −98 and −126 CpG sites of *STING1*, while only depletion of EHMT2 or UHRF1 led to decreased methylation of CpG sites at −220 (Fig. 5j). These results are consistent with reports that EHMT2-UHRF1-DNMT1 promotes genomic DNA methylation^31, 32^. Additionally, our results strongly suggest that the −220, and perhaps −98, CpG site is key for the regulation of *STING1* expression.

### EHMT2 inhibition enhances the efficacy of anti-PD-1 immunotherapy

As increased intratumoral lymphocytes infiltration and PD-L1 expression are predictors for good response to anti-PD-1/PD-L1 based ICIs therapy^33, 34^, we tested whether a UNC0642+anti-PD-1 combination therapy could achieve optimal tumor control. We treated 4T1 tumor bearing BALB/c mice with vehicle+isotype IgG, UNC0642+isotype IgG, vehicle+anti-PD-1, and UNC0642+anti-PD-1 and monitored tumor growth (Fig 6a). UNC0642 at the dose used (10 mg/kg every two days) showed no effect on tumor growth, and anti-PD-1 (200 μg per mouse every two days) alone also failed to produce any therapeutic effect, consistent with published studies^28, 35^. In stark contrast, tumor growth was strongly inhibited in all 6 mice treated with the combination therapy, with some tumors showing clear shrinkage (Fig. 6b, c, d). IHC analyses again showed high STING expression and CD4+ and CD8+ T lymphocytes infiltration in UNC0642 treated tumors (Extended Data Fig. 6a and b). Reduced proliferation (Ki67), and increased apoptosis (cleaved caspase3) were only observed in UNC0642+anti-PD-1 treated tumors (Fig. 6e). Taken together, these data demonstrate that EHMT2 inhibition can substantially enhance the antitumor efficacy of anti-PD-1 and may even overcome the resistance to this therapy.

**Figure 6.**
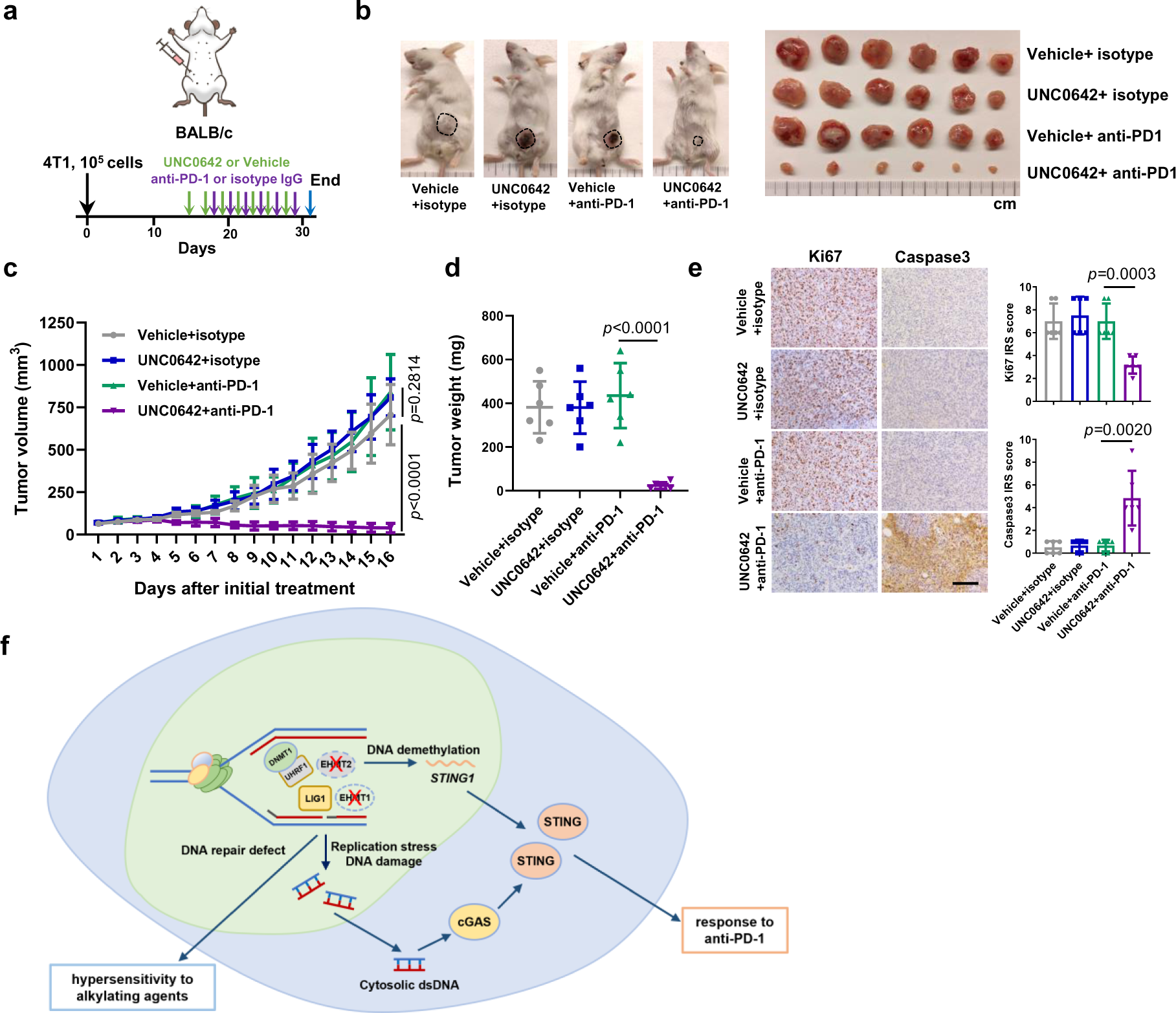
EHMT2 inhibition enhances antitumor response of anti-PD-1 treatment in 4T1 syngeneic allograft tumors. **a** Schematic diagram of 4T1 allograft models and experimental timeline. **b** Representative images of 4T1 tumor bearing mice (left) and an image of all the mammary tumors harvested at study endpoint (day 29). **c** Growth curves of 4T1 tumors from the first day of treatment (day 14) to study endpoint (day 29). **d** Weight of 4T1 tumors at study endpoint (day 29). **e** Representative H&E and IHC images of 4T1 tumors (left) and quantification of the IHC results using the immunoreactive score (IRS, right). Scale bar, 50 µm. **f** Model of EHTM1/EHMT2 inhibition induced hypersensitivity to the alkylating agents or immune checkpoint inhibitors. Data (**c**, n=6; **d**, **e**, dots=6) are presented as mean ± standard deviation (SD). Statistical test is preformed using two-tailed unpaired Student’s *t* test and the exact *p* value is provided.

## Discussion

In this study, we show that EHMT inhibition causes slowdown of replication fork progression, accumulation of ssDNA replication gaps, and cellular sensitivity to alkylating agents. Mechanistically, the above effects of EHMT inhibitors result primarily from the inhibition of LIG1 methylation by EHMT1 (and not EHMT2). In addition to the reported role of enabling LIG1 interaction with UHRF1, we find that EHMT1-mediated methylation of LIG1 also promotes its association with chromatin to guarantee normal DNA replication and repair of alkylated DNA. On the other hand, we demonstrate that EHMT2-UHFR1 deficiency activates the cGAS-STING pathway, at least in part, by *STING1* promoter demethylation and transcriptional depression, while EHMT1-LIG1 deficiency is unable to elicit such an effect (Fig. 6f). Together, our results highlight the distinct function of these two paralogs that are commonly thought to function together as a heterodimeric complex.

Data on the impact of LIG1 status on cancer therapy response has been controversial. It has been reported that LIG1 deficient ovarian cancer cells show higher cisplatin sensitivity compared to proficient cells^36^, however another study reported that loss of LIG1 in patient derived TNBC xenografts correlates to carboplatin resistance^37^. Consistent with early findings^38, 39^, our results show that LIG1 deficiency particularly sensitizes cancer cells to alkylating agents, suggesting that alkylating agents may be effective for cancers with LIG1 deficiency. Importantly, we also demonstrate that UNC0642 strongly potentiates the efficacy of CTX, a commonly used chemotherapy for the treatment of breast and several other cancers, in two different mouse models of TNBC by compromising the function of WT LIG1, raising the prospect that combining alkylating agents with EHMT1 inhibition may be efficacious for cancers with normal LIG1 function, including poor prognosis tumors such as TNBC and GBM.

In recent years, replication stress has gained increasing recognition as both a key feature and a targetable vulnerability of cancer cells. In our present study, EHMT1 inhibition and CTX treatment each leads to replication stress as revealed by reduced fork progression and formation of replication gaps, and their combination elicits even stronger replication stress accompanied by extensive ssDNA regions that presumably include both replication gaps and repair gaps. Our results suggest that in the setting of LIG1 loss or EHMT1 inhibition, the MRE11 nuclease processes the unligated Okazaki fragments or DNA repair intermediates into extensive ssDNA gaps, which may cause RPA exhaustion and, perhaps together with increased DSBs, trigger the activation of cell death pathways leading to enhanced cancer cell killing (Extended Data Fig. 4). Therefore, our results are consistent with the notion that ssDNA gaps are a key factor in therapy response^40^, and the ability of mice to tolerate the UNC0642-CTX combination while the tumor growth was severely affected also suggests that certain tumor types, including TNBC, may be more sensitive to ssDNA gaps than normal cells.

The cGAS-STING pathway, typically activated by cytosolic DNA, has been recognized as a key determinant of tumor immune microenvironment and response to immunotherapy^41^. In the present study, EHMT inhibitors induce STING expression in tumor cells leading to chemocytokine production, T cells infiltration and remarkable response to anti-PD-1 immunotherapy. Notably, the induction of STING is at the mRNA level, and depletion of EHMT2, and not EHMT1, leads to increased STING expression, suggesting that the mechanism may be independent of the DNA damage induced by EHMT inhibition. Interestingly, depletion of UHRF1, and not LIG1, also induces STING mRNA expression. Given that EHMT2 and UHRF1 have been reported to either bind each other or co-exist in certain complexes^31, 42, 43^, our findings point to an EHMT2-UHRF1-DNMT1 axis that suppresses STING promoter activity, presumably by methylating the CpG sites therein, especially the ones at −220 and −98 nt relative to its TSS. The fact that EHMTi also induced STING mRNA expression indicates that EHMT2 enzymatic activity is involved, and the precise mechanism awaits further investigation. Whether the cGAS promoter is under similar regulation also remains to be seen.

The effect of immune checkpoint inhibitors (ICIs) therapy on patients with TNBC has been debated as only a limited portion of clinical cases receive long term benefits from this therapy^44, 45^. Mechanisms of insensitivity, or “resistance”, to ICIs include defects in antigen expression in tumor cells, decreased chemocytokines release, insufficient effective T cells intratumoral infiltration, and so on^46, 47^. Activation of cGAS-STING pathway has emerged as a key determinator and predictor of ICIs therapy response^48–50^. However, STING expression is generally low in breast and other cancers showing an immunosuppressed phenotype^51, 52^. Therefore, pharmacologic induction of STING expression is a logical approach to engineer response to ICIs therapy. In this vein, EHMT inhibitors induce both STING expression and cytosolic DNA, which activates the cGAS-STING pathway, therefore representing an ideal avenue to improve the response rate and efficacy of ICIs therapy. Additionally, EHMTi-induced DNA demethylation may also promote antigen expression in tumor cells potentially converting poorly immunogenic tumors to “hot” tumors, which may further increase the anti-tumor effects of intratumoral immune cells.

## Acknowledgements

We thank Dr. Yibin Kang for providing the MDA-MB-231-luc cells. This study was supported by the US National Cancer Institute (R01CA138804, R01CA262227, and P01CA250957-9485 to BX, P01CA250957-9486, R01CA243547, and P30-CA072720-5928 to SG). ZK was supported by a postdoctoral fellowship from the New Jersey Commission of Cancer Research (NJCCR, DCHS20PPC005).

## Author contributions

BX and ZK conceived the project. ZK performed most of the experiments and data analyses. PF performed part of DNA fiber assays, drug sensitivity and data analysis. HM, TL performed part of cell and mouse work. KL performed part of plasmids construction and cell cycle analysis. TL performed IHC. JL performed IVIS imaging. VG and PR provided reagents and expertise. ZF, MG and SG served as consultants. ZK drafted the manuscript. SG edited the manuscript. BX supervised the study and edited the manuscript.

## Materials and methods

### Cell lines

U2OS, MDA-MB-231 and HEK293T cells were purchased from ATCC and cultured in Dulbecco’s modified Eagle’s medium (DMEM) (Sigma, 5796). MDA-MB-231-luc cells were provided by Dr. Yibin Kang (Princeton University) and also grown in DMEM. 4T1 cells were also purchased from ATCC and cultured in RPMI1640 (Sigma, R8758). Both media were supplemented with 10% fetal bovine serum (Sigma, F0926) and 1% penicillin and streptomycin (Sigma, P0781). All cell lines were cultured at 37°C in a humidified incubator with 5% CO2.

### Plasmid constructs

The LIG1 open reading frame (ORF) was amplified by PCR from pDONR223_LIG1_WT_V5 (Addgene, 83006) and cloned into the pOZ-FH-C vector. To allow for selective depletion of endogenous LIG1 in cells harboring this construct, 4 silent base changes were introduced into the binding site of LIG1 siRNA 1659 by site-directed mutagenesis, using the QuikChange II XL Site-directed mutagenesis kit (Agilent Technologies, 200521). After sequencing confirmation, site-directed mutagenesis was again conducted to generate LIG1 K126R and K126A mutants. Details will be provided upon request.

### RNA interference

siRNAs targeting EHMT1, EHMT2, LIG1 and UHRF1 were custom synthesized by Sigma. AllStars negative control siRNA was purchased from Qiagen (1027281). The sense strand sequences of the siRNAs are listed in Supplementary Table 1. Transfections were performed using Lipofectamine RNAiMax (Thermo Fisher, 13778150) following manufacturer’s instructions. The final concentration of siRNAs was 8 nmol/L.

### Cell lysates, cell fractionation, immunoprecipitation (IP) and western blotting

Cells were harvested by trypsinization and washed with PBS. For generation of whole cell homogenates, cells were lysed in NETNG350 (20 mM Tris-HCl [pH 7.5], 350 mM NaCl, 1 mM EDTA, 10 mM NaF, 0.5% NP-40, and 5% glycerol) lysis buffer on ice for 10 min, and the mixtures were then sonicated with a Fisher Model 100 Sonic Dismembrator at setting 2 for 5 rounds (10 sec in total). For fractionation, cells were lysed with NETNG100 (100 mM NaCl, otherwise the same as NETNG350) on ice for 10 min, the lysates were centrifuged at 15,000 x g for 10 min, and the supernatants were collected as the soluble fractions. The pellets, considered as chromatin fractions, were re-suspended in NETNG350 and sonicated as above. For IP, cells were lysed in NETNG250, and IP was carried out using anti-FLAG M2 beads (Sigma, A2220) to pull down FLAG-HA-tagged LIG1 on a rocker at 4°C for overnight. All lysis buffers were added with the Complete protease inhibitor cocktail (Roche, 11697498001).

Protein samples were electrophoresed on home-made 4-16% Tris-glycine SDS-polyacrylamide gels and transferred to nitrocellulose membranes (0.45 µm, Bio-Rad, 1620115). Membranes were blocked in 5% non-fat milk, incubated with primary and secondary antibodies, and developed using Immobilon Western Chemiluminescent HRP Substrate (Millipore, WBKLS0500). All experiments were repeated 3 times independently. Antibodies used are listed in Supplementary Table 2.

### BrdU incorporation and flow cytometry

U2OS cells were plated into 6-well plates and treated with 4 µM UNC0638 or UNC0642 for 54 h. Cells were labeled with 100 µM BrdU for 20 min in pre-wormed fresh media, trypsinized and transferred to Eppendorf tubes. Cells were washed with PBS with 1% BSA and then fixed in 70% ethanol at −20°C overnight. To denature the DNA, cells were spun down and resuspended in 2 M HCl with 0.5% Triton X-100 at room temperature for 30 min. After centrifugation, cells were re-suspended in 0.1 M Na_2_B_4_O_7_ (pH 8.5) to neutralize the acid. After washing with 1% BSA/PBS, cells were resuspended in 300 µL PBS with 1% BSA and 0.5% Tween 20, 20 µL FITC anti-BrdU (BD Pharmingen, 347583) was then added, and the mixture was incubated on a rocker at 4°C overnight. Cells were then washed with the above BSA/Tween 20/PBS, re-suspended in PBS containing 1 µg/mL propidium iodide (PI), and analyzed on an Attune flow cytometer (Thermo Fisher). All experiments were repeated 3 times independently. Drug information is listed in Supplementary Table 3.

### DNA Fiber Assay

Active replication forks in living cells were labeled by sequential pulses of two thymidine analogs, CldU (20 μM) followed by IdU (200 μM). For U2OS and MDA-MB-231 cells, each pulse was for 20 min, and for 4T1 cells, which grow much faster, each pulse was for 15 min. Cells were then collected by trypsinization, washed with PBS, and re-suspended in cold PBS. For assays involving S1 nuclease treatment, after IdU pulse, cells were incubated in CSK100 buffer (100 mM NaCl, 3 mM MgCl2 [pH 7.2], 300 mM sucrose, 10 mM MOPS, and 0.5% Triton X-100) for 10 min at room temperature, followed by incubation in S1 nuclease buffer with or without 20 U/mL S1 nuclease (Thermo Fisher, 18001016) for 15 min at 37°C. Cells were then scraped and collected in 0.1% BSA/PBS, centrifuged at 7,000 rpm for 5 min at 4°C, and re-suspended in cold PBS.

A total of 3×10^3^ cells in 2 μL of cell suspension were spotted on one end of a glass slide and lysed by 7.5 μL of lysis solution (50 mM EDTA and 0.5% SDS in 200 mM Tris-HCl [pH 7.5]) at room temperature. DNA molecules were spread along the slides by tilting at approximately 15 degrees and then fixed in methanol/acetic acid (3:1) at −20°C for 15 min. After washing with water, the slides were incubated in 2.5 M HCl for 80 min to denature DNA, washed with PBS, and then blocked with blocking buffer (5% BSA/PBS) for 20 min. Slides were incubated with primary antibodies at 4°C overnight and secondary antibodies at 37°C for 1.5 h (both in blocking buffer). Slides were washed with PBS and mounted with ProLong Gold Antifade Mountant (Thermo Fisher, P36930). Images were acquired with a Nikon TE2000 fluorescence microscope, and at least 200 individual IdU labeled tracts from these images were measured using ImageJ 1.51j8 software. Antibodies used are listed in Supplementary Table 2.

### Generation of U2OS cell lines stably expressing LIG1 proteins

Cell lines were generated by transducing U2OS cells with retroviruses expressing C-terminally FLGA-HA-tagged LIG1 WT or K126 mutant proteins. Viruses were packaged by co-transfecting HEK293T cells with pOZ-FH-C-LIG1, gag-pol and env plasmids using X-tremeGENE HP transfection reagent (Sigma, 6366236001). The supernatants were used to infect U2OS cells, and infected cells were selected with Dynabeads Goat Anti-Mouse IgG (Invitrogen, 11033) coupled with anti-IL-2 Rα antibody (Sigma, clone 7G7/B6).

### Drug sensitivity assay

To assess the effect of LIG1, EHMT1 or EHMT2 depletion on drug sensitivity, U2OS or MDA-MB-231 cells were transfected with siRNAs for 30 h, trypsinized and reseeded into 96-well plates at 2,000 cells per well, and incubated overnight. Cells were then treated with drugs at different dosages. To test the effect of EHMT1/2 inhibition, cells were seeded into 96-well plates at 2,000 cells per well for MDA-MB-231 and U2OS cells, or 600 cells per well for 4T1 cells, and allowed to adapt overnight. Cells were then treated with a fixed concentration of UNC0638 or UNC0642 (2 µM for both) in combination with chemotherapeutic drugs at different dosages. Cell viability was measured 72 h after drug treatment using CellTiter-Glo 2.0 Cell Viability Assay (Promega, G9241) following manufacturer’s instructions.

### BrdU and RPA32 immunofluorescence (IF)

MDA-MB-231 cells were seeded onto coverslips and subjected to siRNA knockdown or drug treatments for various periods of time. Cells were then labeled with BrdU (20 μM) for 24 h, followed by treatment with mirin (50 μM for 2 h) and/or CTX (9 mM for 6 h) where applicable. Briefly, coverslips were washed with PBS, incubated sequentially with extraction buffer 1 (10 mM PIPES [pH 7.0], 100 mM NaCl, 300 mM Sucrose, 3 mM MgCl_2_, 1 mM EGTA, 0.5% Triton X-100) and extraction buffer 2 (10 mM Tris-HCl [pH 7.5], 10 mM NaCl, 3 mM MgCl_2_, 1% Tween 40, 0.5% sodium deoxycholate) for 10 min each. Coverslips were then fixed with 3% paraformaldehyde (PFA), permeabilized with 0.5% Triton X-100/PBS, blocked with 5% BSA/PBS, incubated with primary antibodies overnight at 4°C followed by secondary antibodies at 37°C for 1.5 h, and mounted with ProLong Gold Antifade Mountant (Thermo Fisher, P36931). Images were acquired and 50 cells were analyzed. All experiments were repeated 3 times independently. Antibodies used are listed in Supplementary Table 2.

### Animal studies

Normal and nude BALB/c mice were purchased from Charles River. For MDA-MB-231 xenografts, 3×10^6^ MDA-MB-231-luc cells were suspended in 100 µL of PBS and injected into the mammary fat pads of 5 to 6 weeks old female BALB/c nude mice. Tumor size (long diameter and short diameter) was measured every day with a caliper and tumor volume calculated as 0.5×length×width×width. For chemotherapy, when the tumor volume reached 60-100 mm^3^, mice were separated into 4 groups and each treated with vehicle (55% PBS, 40% PEG300, 5% Tween 80) (same volume of DMSO compared to UNC0642 were added), UNC0642 (5 mg/kg), CTX (20 mg/kg) (same volume of DMSO compared to UNC0642 were added), or a combination of UNC0642 and CTX (5 mg/kg and 20 mg/kg, respectively), via intraperitoneal (i.p.) injection each day for 7 consecutive days.

For 4T1 allografts, 10^5^ 4T1 cells were inoculated into the mammary fat pads of 7 to 8 weeks old female normal BALB/c mice. Tumor size measurement and chemotherpay treatment were performed in the same way as above. For anti-PD-1 treatment, when the 4T1 tumor volume reached 60-100 mm^3^, mice were separated into 4 groups: Vehicle+isotype, UNC0642+isotype, Vehicle+anti-PD-1 and UNC0642+anti-PD-1. Vehicle+isotype and Vehicle+anti-PD-1 mice were treated with vehicle (55% PBS, 40% PEG300, 5% Tween 80, plus the same volume of DMSO as UNC0642), and UNC0642+isotype and UNC0642+anti-PD-1 mice were treated with UNC0642 (10 mg/kg) via intraperitoneal (i.p.) injection every two days. Vehicle+isotype and UNC0642+isotype mice were treated with 200 μg rat IgG2a isotype control (BioXCell, BE0089) in PBS, and Vehicle+anti-PD-1 and UNC0642+anti-PD-1 mice were treated with 200 μg anti-PD-1 (BioXCell, BE0273) via intraperitoneal (i.p.) injection every two days. Study endpoint was when the largest tumor reaching 1,000 mm^3^, at which point mice were euthanized and tumors harvested. Prior to collection of MDA-MB-231-luc tumors, mice were injected with 100 µL of 15 mg/mL D-Luciferin (Perkin Elmer, LLC 122799), anesthetized in an induction chamber using Isoflurane gas, and imaged using the IVIS Lumina LT In Vivo Imaging System (Perkin Elmer, CLS136331) before being euthanized. All animal works were conducted under a protocol (ID999900493) approved by the Institutional Animal Care and Use Committee (IACUC) of Rutgers University.

### Histology

Mouse tumor tissues were fixed in 10% formalin for 48 h, embedded in paraffin and cut into 5-µm sections, then H&E staining and immunohistochemistry (IHC) were conducted. IHC results were scored based on the percentage of positive cells and intensity of staining. Namely, the immunoreactive score (IRS) were calculated as A (percentage of positive cells, 0= no positive cells, 1= <10%, 2= 10-50%, 3= 51-80% or 4= >80%) × B (0= no color reaction, 1= mild reaction, 2= moderate reaction, 3= intense reaction). Antibodies used for IHC are listed in Supplementary Table 2.

### Quantitative real time PCR (qRT-PCR) analysis

Total RNA was isolated from U2OS and MD-MB-231 cells using the RNeasy Mini Kit (QIAGEN no. 74134). cDNA was synthesized using Superscript III First-strand Synthesis System for RT-PCR kit (Invitrogen, 18080-051) following the manufacturer’s instructions. qRT-PCR reactions were performed using PowerUp SYBR Green Master Mix (Thermo Fisher, A25742). PCR reaction was carried out on QuantStudio 3 Real-Time PCR System (Thermo Fisher, A28132) and thermal cycling conditions included 50°C for 2 min and an initial denaturation step of 95°C for 10 min and 40 cycles at 95°C for 15 s and 60°C for 60 s. Analyses were carried out in triplicate for each data point. The PCR primers used for gene expression analysis are listed in Supplementary Table 1.

### Pico488 Staining

U2OS cells were seeded onto coverslips and subjected to DMSO, UNC0638 and UNC0642 treatment (4 μM for 48 h). To detect cytosolic DNA, cells were incubated with cell medium with Pico488 double-stranded DNA quantification reagent (Lumiprobe, 42010) for 1 h, washed with PBS, and then incubated in dye-free medium for 20 min. Coverslips were then rinsed three times in PBS, fixed in 3% paraformaldehyde (PFA) for 15 min, and then mounted with ProLong Gold Antifade Mountant (Thermo Fisher, P36931). Images were acquired with a Nikon TE2000 fluorescence microscope.

### Bisulfite sequencing DNA methylation analysis

Genomic DNA was extracted from U2OS cells by using the DNeasy Blood & Tissue Kit (Qiagen, 69504). The purified DNA was chemically modified with the use of EZ DNA Methylation-Gold Kits (ZYMO RESEARCH, D5005) according to the manufacturer’s instructions, to detect methylated CpG sites. During this modification, all unmethylated cytosines in genome DNA exposure to sodium bisulfite will be converted to uracils, whereas leaving methylated cytosines unaltered.

PCR primers (Supplementary Table1) were designed to amplify a 316 bp region of the *STING1* promoter that contains 6 CpGs (Extended Data Fig. 6a). The PCR product was then subcloned into the pCR4-TOPO vector by using TOPO TA Cloning Kit (Thermo Fisher, K4575J10). Recombinant plasmids from five clones of each sample were sent to Psomagen (Brooklyn, New York) to complete the bisulfite DNA-sequencing procedure.

### Statistical analyses

All experiments were performed at least three times independently. Statistical significances were determined using unpaired or paired Student’s *t* test as appropriate. All statistical analyses were carried out using GraphPad Prism 7.0.

---

**Extended Data Figure 1.**
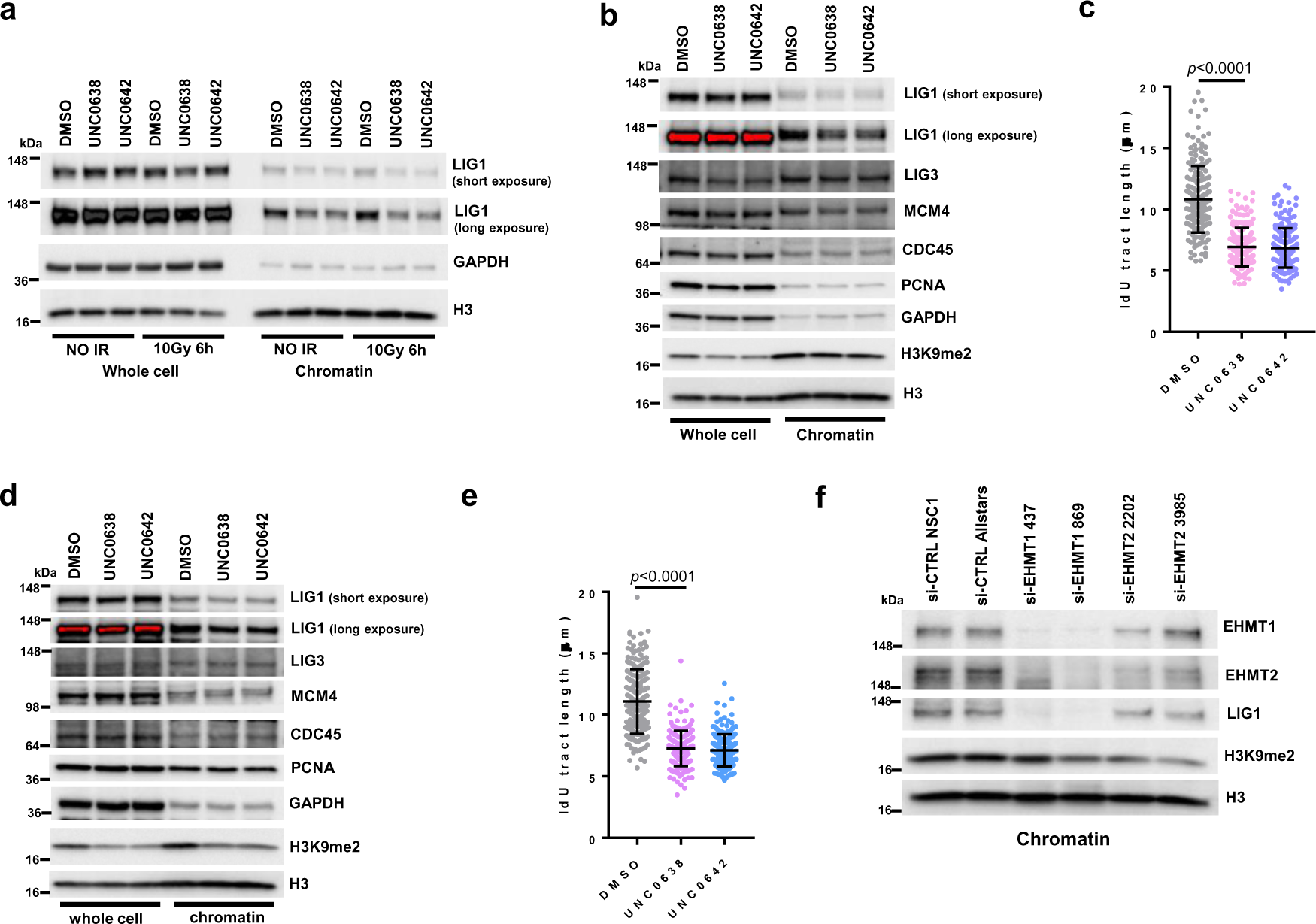
EHMT1/2 inhibition destabilizes LIG1 in chromatin to restrain replication fork progression. **a** Western blots showing LIG1 in U2OS cells treated with UNC0638 and UNC0642 4µM for 54h with and without ionzing radiation 10Gy. **b** Western blots showing DNA replilcation proteins in MDA-MB-231 cells treated with UNC0638 and UNC0642 3µM for 72h. **c** Lengths of IdU labeled replication tracts in MDA-MB-231 cells treated with UNC0638 and UNC0642 3µM for 72h. **d** Western blots showing DNA replilcation proteins in 4T1 cells treated with UNC0638 and UNC0642 3µM for 72h. **e** Lengths of IdU labeled replication tracts in 4T1 cells treated with UNC0638 and UNC0642 3µM for 72h. **f** Western blots showing levels of LIG1 from chromatin fractions in EHMT1- and EHMT2-depleted U2OS cells. Data (**c** and **e**, dots=200) are presented as mean ± standard deviations (SD). Statistical test is preformed using two-tailed unpaired Student’s *t* test and the exact *p* value is provided.

**Extended Data Figure 2.**
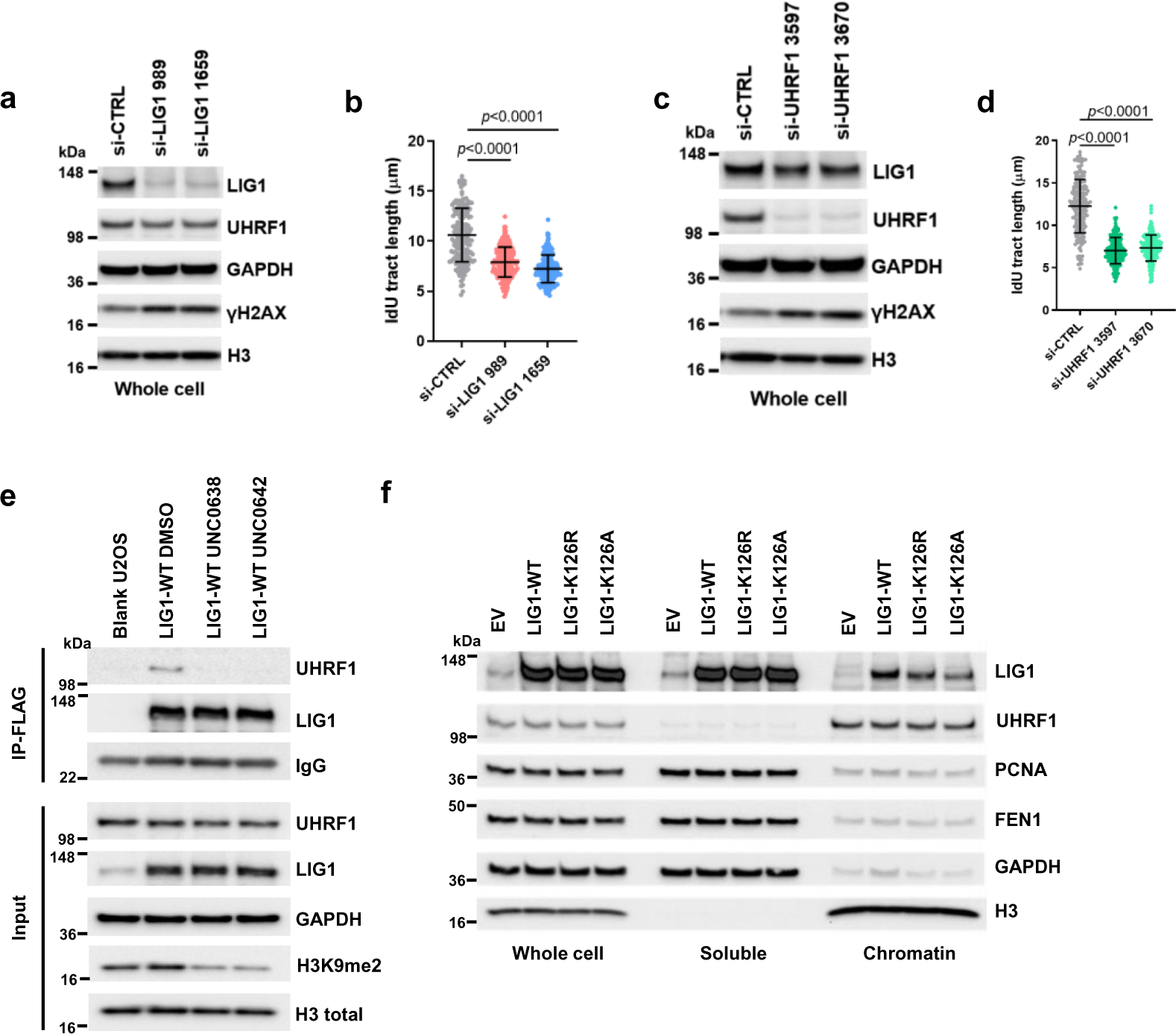
Disruption of LIG1-UHRF1 binding destabilizes LIG1 in chromatin. **a** Protein levels in LIG1-depleted U2OS cells.. **b** Lengths of IdU labeled replication tracts in LIG1-depleted U2OS cells. **c** Protein levels in UHRF1-depleted U2OS cells. **d** Lengths of IdU labeled replication tracts in UHRF1-depleted U2OS cells. **e** Co-IP of endogenous UHRF1 with FLAG-HA-tagged WT LIG1 in U2OS cells stably expressing the protein treated with DMSO, UNC0638 or UNC0642 (4 µM) for 54 h. **f** Protein levels of LIG1, UHRF1, PCNA and FEN1 in the whole cell as well as soluble and chromatin fractions of 293T cells transiently transfected with LIG1 WT, K126R and K126A cDNA constructs. Data (**b** and **d** dots=200) are presented as mean ± standard deviations (SD). Statistical test is preformed using two-tailed unpaired Student’s *t* test and the exact *p* value is provided.

**Extended Data Figure 3.**
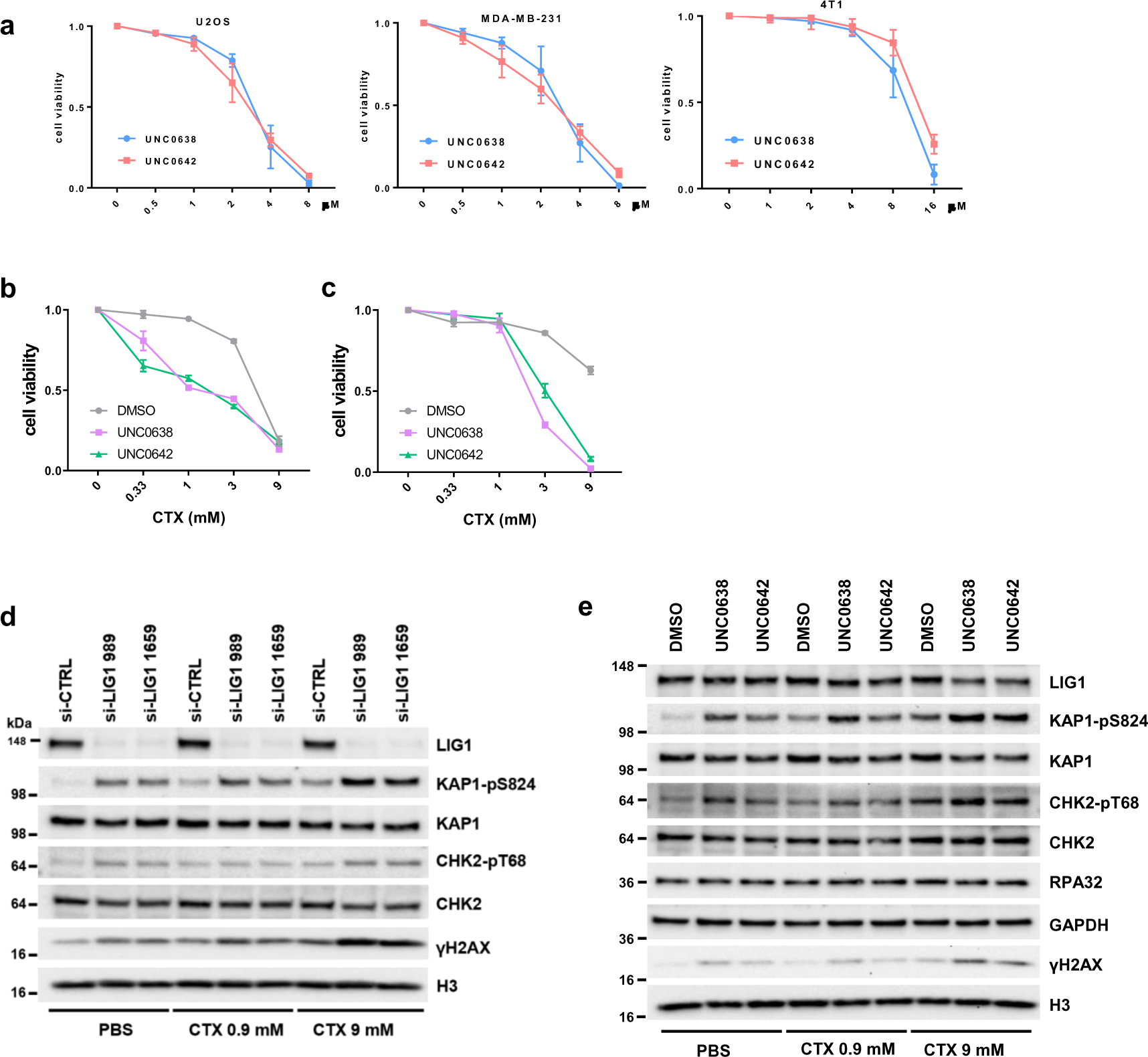
EHMT1 inhibition sensitizes cancer cells to cyclophosphamide (CTX). **a** Sensitivity of U2OS, MDA-MB-231 and 4T1 cells to UNC0638 and UNC0642 for 72h. **b** Survival curves of U2OS cells treated with different concentrations of CTX in combination with a fixed concentration (2 µM) of UNC0638 or UNC0642 for 72 h. **c** Survival curves of 4T1 cells treated with different concentrations of CTX in combination with a fixed concentration (4 µM) of UNC0638 or UNC0642 for 72 h. **d** Western blots showing levels of KAP1, CHK2 and H2AX phosphorylation in MDA-MB-231 cells treated first with LIG1 depletion and then with CTX (9 mM for 24 h). **e** Western blots showing levels of KAP1, CHK2 and H2AX phosphorylation in MDA-MB-231 cells treated first with UNC0638 or UNC0642 (4 µM for 30 h) and then with CTX (9 mM for 24 h).

**Extended Data Figure 4.**
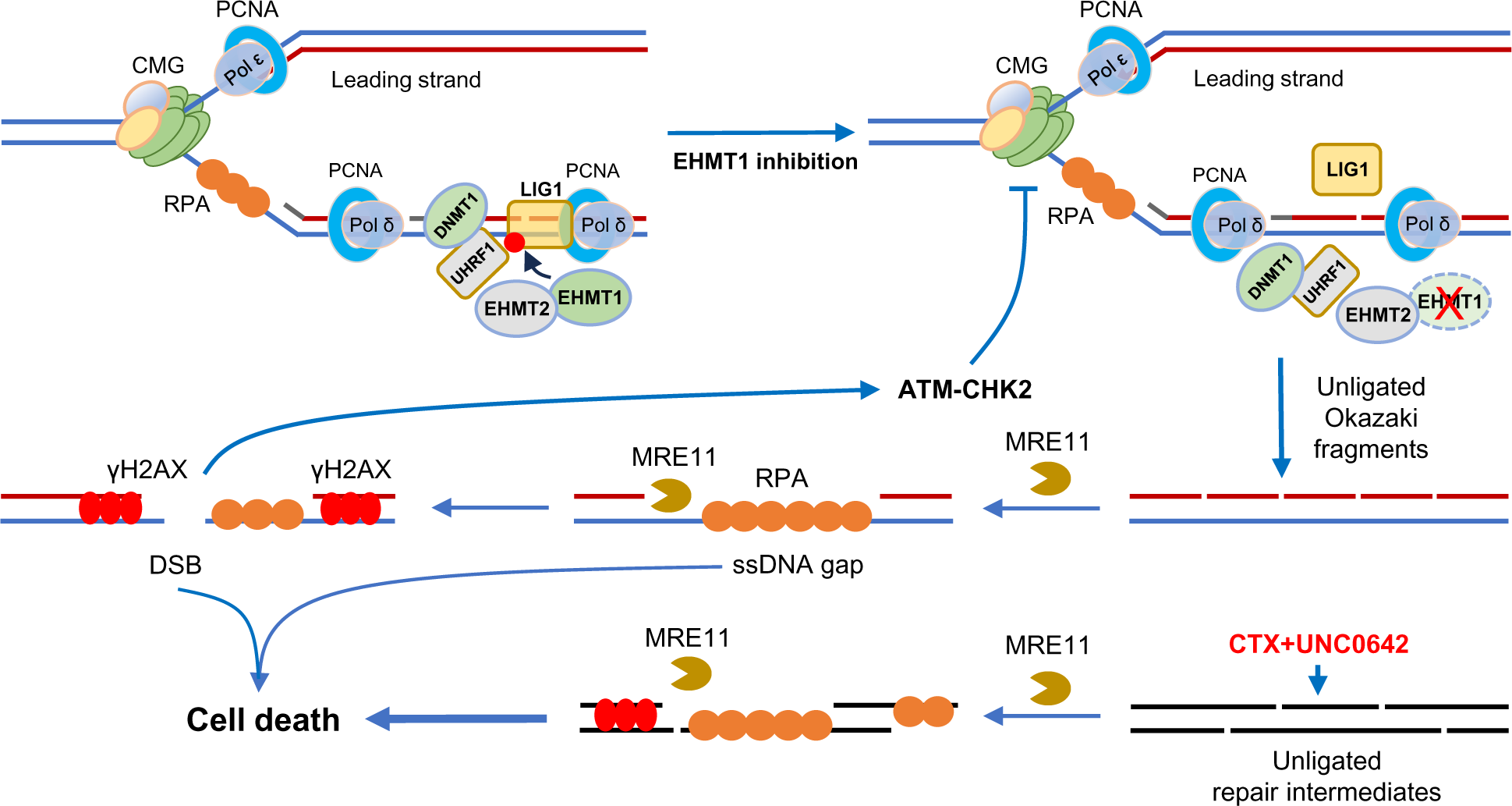
A proposed model of EHMT1 inhibition-induced replication stress and sensitization to alkylating agents. Under normal conditions, EHMT1 methylates LIG1, which promotes its chromatin association and interaction with UHRF1, thereby maintaining replication fork progression and genome methylation. Upon EHMT1 inhibition, loss of LIG1 methylation results in its reduced association with chromatin and disengagement with UHRF1, leading to unligated Okazaki fragments at the replication fork, generation of ssDNA gaps by MRE11, DNA DSBs, and activation of ATM, which restrains replication fork progression. During CTX+EHMT1 combination therapy, CTX generates adducts on DNA, whose repair by NER and BER requires LIG1 for final ligation of nicks on DNA; EHMT1 inhibition compromise LIG1 chromatin recruitment and its complex formation with UHRF1, which together lead to accumulation of unligated nicks and unrepaired SSBs on DNA throughout the genome, including replication forks. MRE11 aberrantly processes the nicks and SSBs into extensive and long stretches of ssDNA, leading to RPA exhaustion and DNA breakage, triggering cell death pathways to execute cell killing.

**Extended Data Figure 5.**
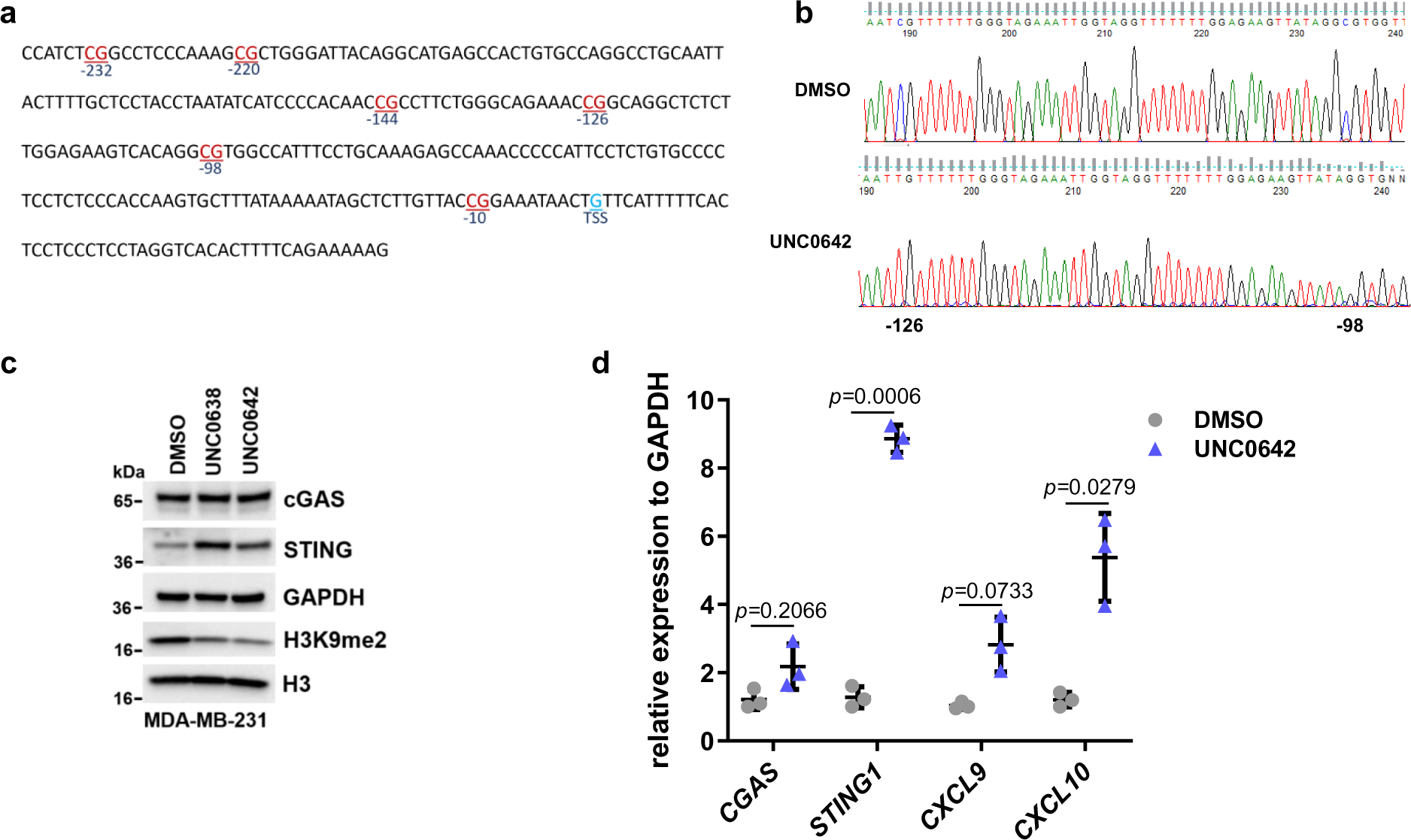
EHMT2 inhibition promotes STING expression through demethylation of *STING1* promoter. **a** The CpG sites in a −240nt region upstream of the transcriptional start site (TSS) of the *STING1* promoter. **b** Sequencing of bisulfite converted DNA from U2OS cells treated with UNC0638/0642 (4 µM for 48 h) was performed to determine the percent methylation for each CpG site. **c** Protein levels of STING pathway factors in MDA-MB-231 cells treated with DMSO, UNC0638 or UNC0642 (4 µM for 24 and 48 h). **d** Quantitative real time (qRT)-PCR analysis (n=3) mRNA expression levels of cGAS-STING pathway genes in MDA-MB-231 cells treated with DMSO and UNC0642 (4 µM for 48 h). Data (**d**, dots=3) are presented as mean ± standard deviation (SD). Statistical test is preformed using two-tailed paired Student’s *t* test and the exact *p* value is provided.

**Extended Data Figure 6.**
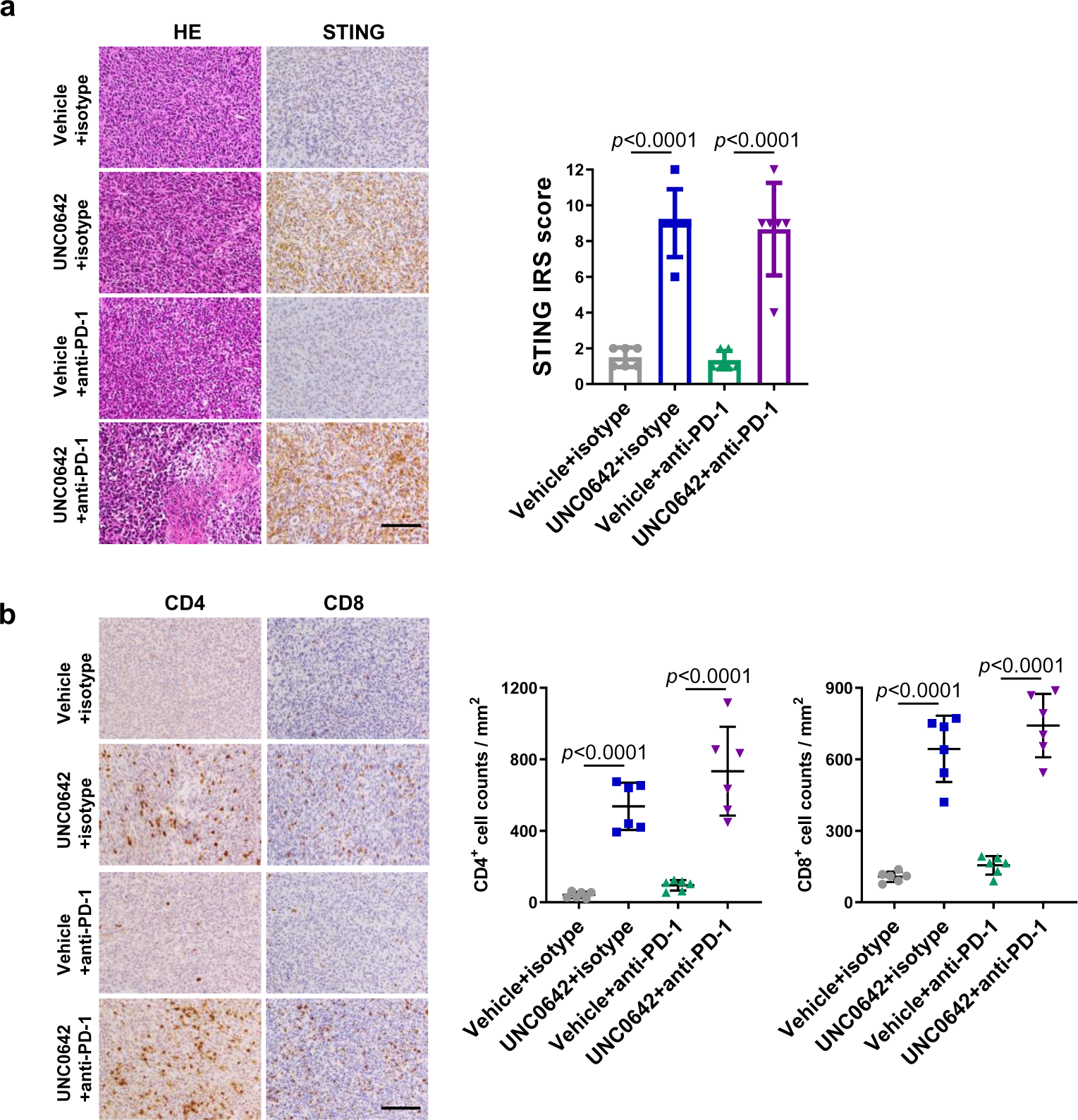
EHMT2 inhibition promotes STING expression and lymphocytes infiltration in 4T1 tumors. **a** Representative HE and IHC images of 4T1 tumors (left) and quantification of the STING expression using the immunoreactive score (right). Scale bar, 50 µm. **b** Representative CD4+ and CD8+ staining IHC images of 4T1 tumors (left) and quantification of the infiltrating lymphocytes (right). Scale bar, 50 µm. Data (**a** and **b**, dots=6) are presented as mean ± standard deviation (SD). Statistical test is preformed using two-tailed unpaired Student’s *t* test and the exact *p* value is provided.

**Supplementary Table 1.**
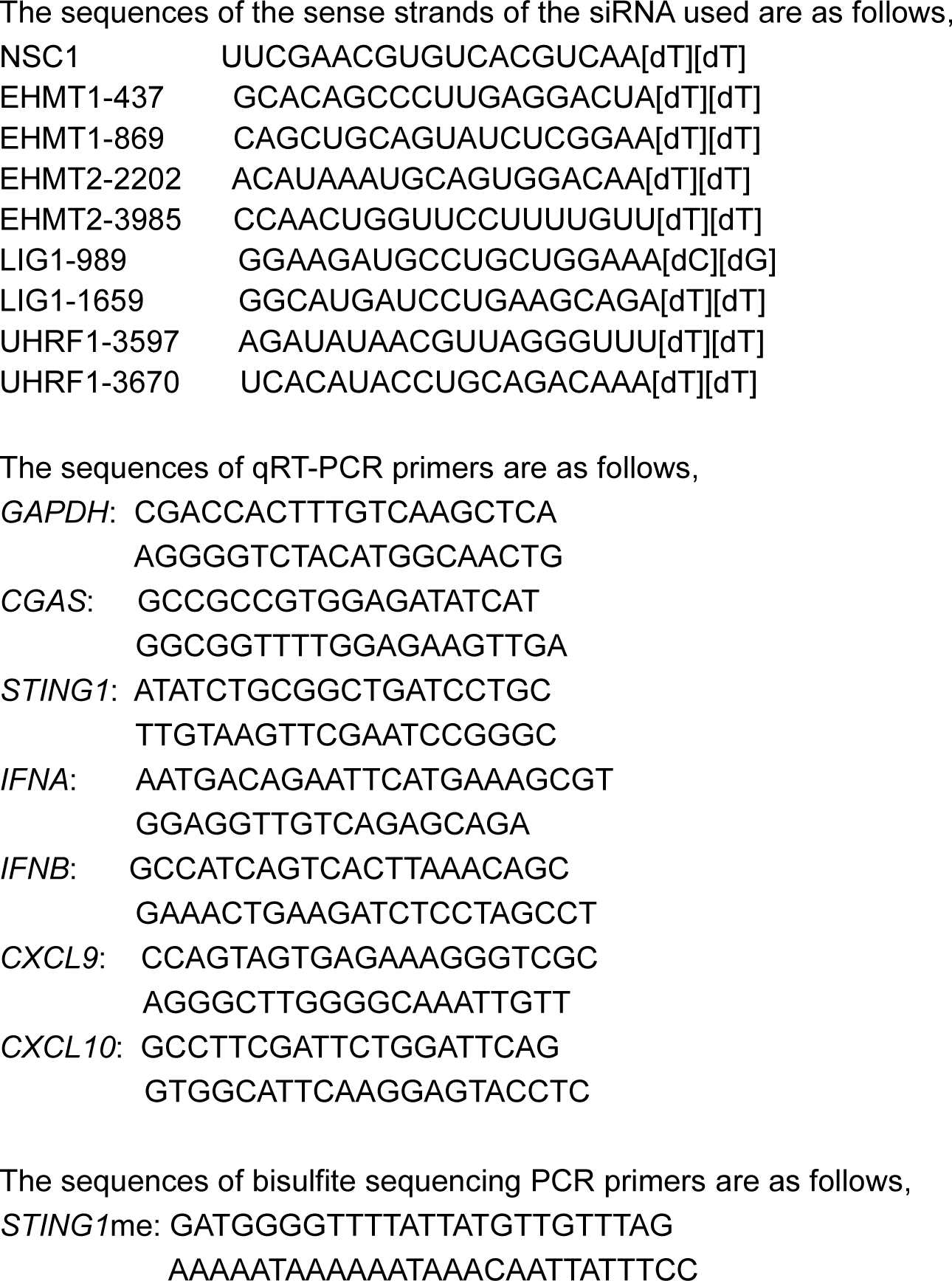
Sequences of the siRNAs and primers in this study.

**Supplementary Table 2.**
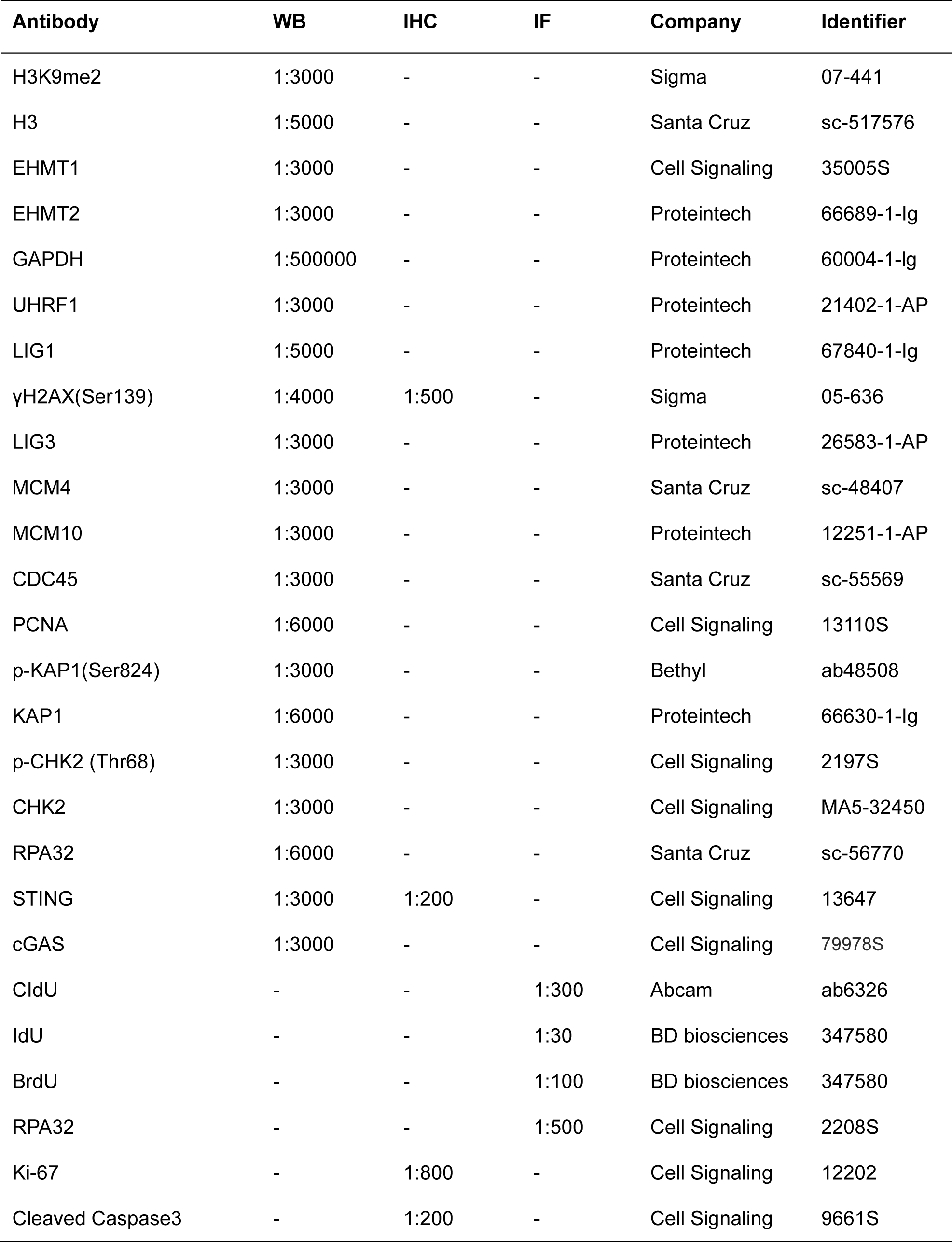
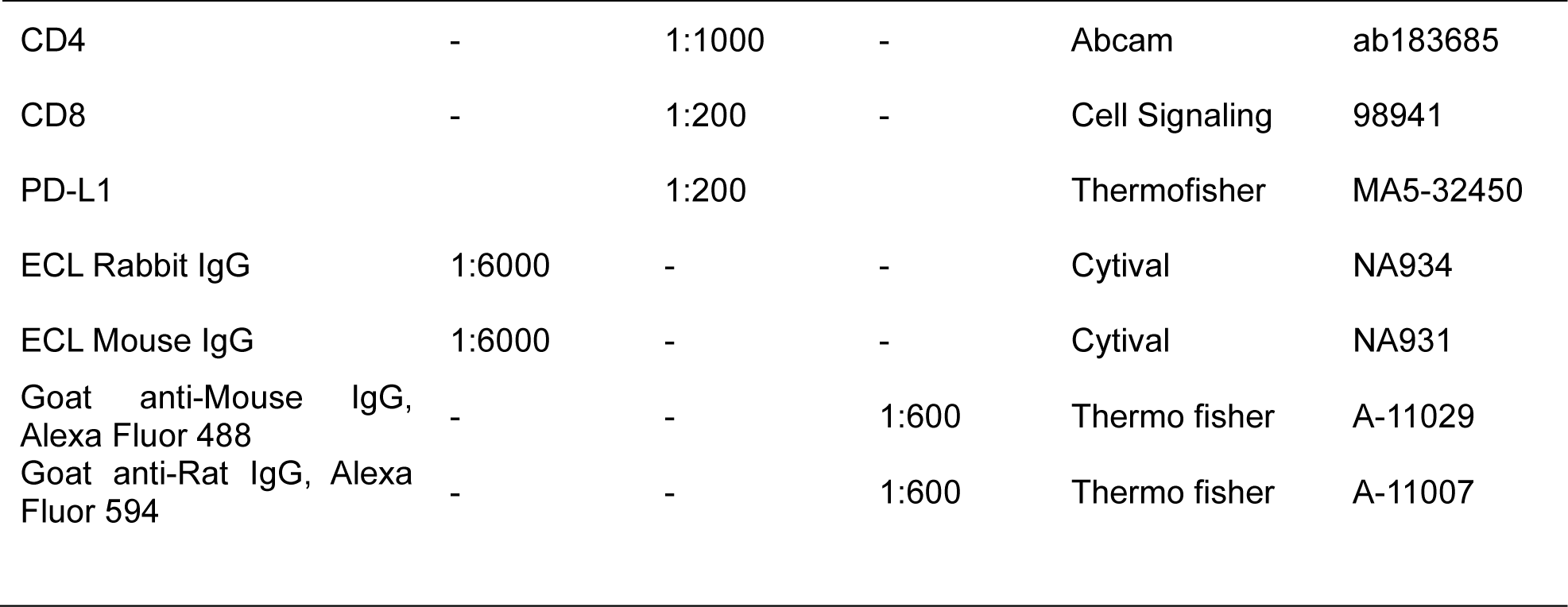
Antibodies for WB, IHC and IF used in this study.

**Supplementary Table 3.** Drugs information used in this study.

| **Drugs** | **Company** | **Identifier** |
| --- | --- | --- |
| UNC0638 | Selleckchem | S8071 |
| UNC0642 | Selleckchem | S7230 |
| BrdU | Sigma | B5002 |
| CldU | Sigma | C6891 |
| IdU | Sigma | I7125 |
| ATMi KU55933 | Selleckchem | S1092 |
| ATMi KU60019 | Selleckchem | S1570 |
| ATRi VE821 | Selleckchem | S8007 |
| Bleomycin | vwr | 1076308 |
| Camptothecin | Selleckchem | S1288 |
| Cisplatin | Sigma | PHR1624 |
| Cyclophosphamide | Selleckchem | S2057 |
| Temozolomide | Sigma | PHR1437 |
| Mirin | Selleckchem | S8096 |
| anti-mouse PD-1 (CD279) | Bioxcell | BE0273 |
| rat IgG2a isotype | Bioxcell | BE0089 |

